# Inferring biochemical reactions and metabolite structures to cope with metabolic pathway drift

**DOI:** 10.1101/462556

**Authors:** Arnaud Belcour, Jean Girard, Méziane Aite, Ludovic Delage, Camille Trottier, Charlotte Marteau, Cédric Leroux, Simon M. Dittami, Pierre Sauleau, Erwan Corre, Jacques Nicolas, Catherine Boyen, Catherine Leblanc, Jonas Collén, Anne Siegel, Gabriel V. Markov

**Affiliations:** Univ Rennes, Inria, CNRS, IRISA, Equipe Dyliss, Rennes, France; Sorbonne Université, CNRS, Integrative Biology of Marine Models (LBI2M), Station Biologique de Roscoff (SBR), 29680 Roscoff, France; LBCM, IUEM, University of Bretagne-Sud, Lorient, France; Sorbonne Université, CNRS, Plateforme METABOMER-Corsaire (FR2424), Station Biologique de Roscoff, Roscoff, France; Sorbonne Université, CNRS, Plateforme ABiMS (FR2424), Station Biologique de Roscoff, Roscoff, France

## Abstract

Inferring genome-scale metabolic networks in emerging model organisms is challenging because of incomplete biochemical knowledge and incomplete conservation of biochemical pathways during evolution. This limits the possibility to automatically transfer knowledge from well-established model organisms. Therefore, specific bioinformatic tools are necessary to infer new biochemical reactions and new metabolic structures that can be checked experimentally. Using an integrative approach combining both genomic and metabolomic data in the red algal model *Chondrus crispus*, we show that, even metabolic pathways considered as conserved, like sterol or mycosporine-like amino acids (MAA) synthesis pathways, undergo substantial turnover. This phenomenon, which we formally define as “metabolic pathway drift”, is consistent with findings from other areas of evolutionary biology, indicating that a given phenotype can be conserved even if the underlying molecular mechanisms are changing. We present a proof of concept with a new methodological approach to formalize the logical reasoning necessary to infer new reactions and new molecular structures, based on previous biochemical knowledge. We use this approach to infer previously unknown reactions in the sterol and MAA pathways.

**Author summary:** Genome-scale metabolic models describe our current understanding of all metabolic pathways occuring in a given organism. For emerging model species, where few biochemical data are available about really occurring enzymatic activities, such metabolic models are mainly based on transferring knowledge from other more studied species, based on the assumption that the same genes have the same function in the compared species. However, integration of metabolomic data into genome-scale metabolic models leads to situations where gaps in pathways cannot be filled by known enzymatic reactions from existing databases. This is due to structural variation in metabolic pathways accross evolutionary time. In such cases, it is necessary to use complementary approaches to infer new reactions and new metabolic intermediates using logical reasoning, based on available partial biochemical knowledge. Here we present a proof of concept that this is feasible and leads to hypotheses that are precise enough to be a starting point for new experimental work.

## Introduction

Reconstruction of genome-scale metabolic networks (GSMs) is a useful and powerful way to integrate data about the metabolism of model organisms due to the increasing availability of genome data [1]. In parallel, metabolomics has maturated as a separate research field, and both are now converging, with the proposal to focus on model organism metabolomes [2]. However, integrating genomic and metabolomic data remains challenging, partly because not all metabolites are indexed in the databases used for GSM reconstruction. For the set of core models prioritized for such approaches, a list of experimentally identified metabolites is not always maintained in a publicly available database [3], and this issue become even more problematic for emerging model species, for which a genome is or will be sooner sequenced, but where the community collecting experimental data is rather limited. Macroalgae belong to this second group of emerging models, which is experiencing drastic changes in research practice due to the availability of high-throughput omics tools [4]. Despite extensive discussion on quality criteria in the field of genome-scale metabolic model reconstruction [5], one missing piece of information is the proportion of metabolites incorporated into the genome-scale metabolic model that are actually described in the literature. Literature data are acknowledged as an important source of knowledge to incorporate into GSMs [6], but current databases tend to point towards bibliographical references concerning pathways or single reactions rather than providing information about the presence of metabolites. A recent survey on 391 metabolites from 21 red, brown, and green macroalgae showed that only 184 of those metabolites were indexed into the PlantCyc database [7]. As a response to this, the metabolomic community is organizing the automation of the taxonomic assignation of metabolites [8].

Integrating data on metabolite presence/absence into GSMs is especially important when working on emerging model organisms that are phylogenetically distant from well-established models, because there are many ways to generate variations during evolutionary time, even within metabolic pathways that may appear to be conserved at first glance. Indeed, even for the well-studied human pathogenic bacterium *Mycobacterium tuberculosis*, high throughput metabolomic screens revealed an unexpected diversity of reactions in central carbon metabolism [9]. Evolutionary models have already been developed to explain the arising of new pathways, with most experimental validations being focused so far at the level of individual enzyme activities [10]. The complementary question, how much conserved pathways remain stable in terms of enzymes, has not yet been addressed in a systematic way. However, very similar issues have been tackled in other subfields of evolutionary biology, and can thus be exported to the field of metabolic pathway evolution.

Developmental system drift has been evidenced some decades ago in the field of animal comparative biology, to explain how morphologically similar structures can be maintained even if there are substantial variations in the molecular mechanisms underlying their formation [11]. The concept was more recently extended to plants, where such cases have been observed in leaf development [12]. It was later exported to the fields of protein evolution [13] and gene expression evolution [14]. We hypothesize that this evolutionary concept also adequately explains the strict conservation of metabolic pathways due to enzymatic replacement by non-orthologous displacement of genes encoding enzymes with identical biochemical function ([15]; Figure 1). A second possible mechanism for metabolic pathway drift, that has the potential to generate observable biochemical diversity in pathways is change in enzyme order, which leads to new biosynthetic intermediates without other changes than their order of intervention (Figure 1). To the best of our knowledge, this second possibility has never been formulated in theory, maybe due to difficulties envisioning an experimental setup to test it.

**Fig 1.**
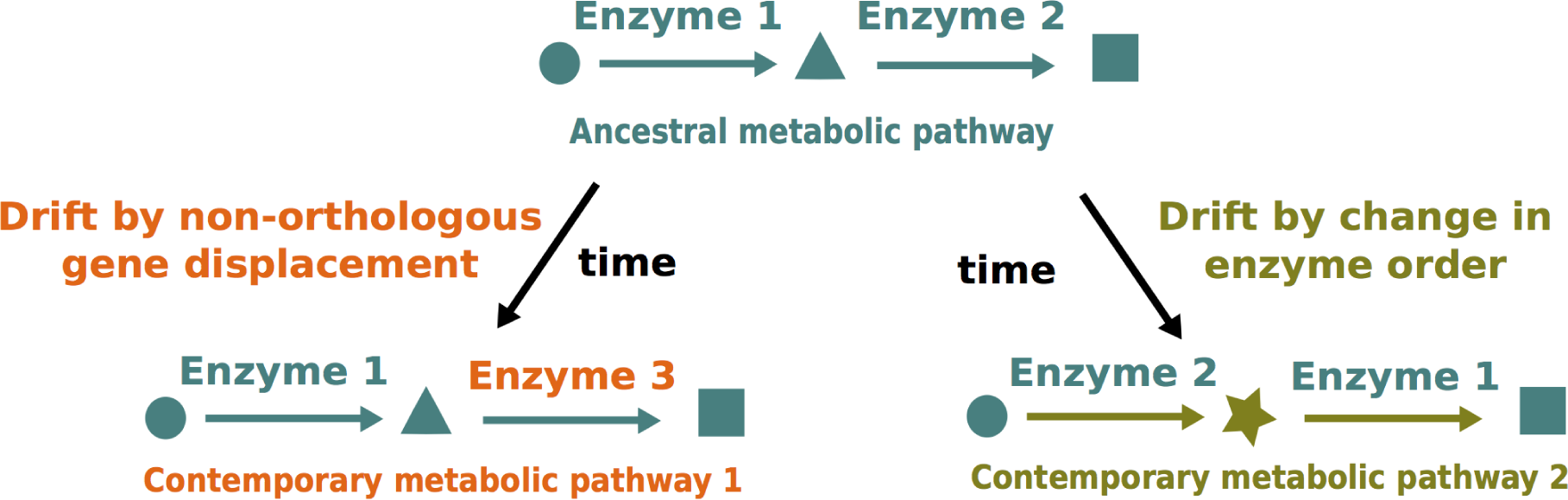
Two possible elementary mechanisms for metabolic pathway drift. Starting from an ancestral pathway (in teal, upper part), changes can occur either by non-orthologous gene displacement (in orange, left side) or change in enzyme order, leading to new metabolites (in olive green, right side).

Regarding non-orthologous gene displacement, classical comparative genomic approaches can generate hypotheses that can be experimentally checked using targeted metabolic profiling combined with enzyme inactivation by CRISPR-Cas9 [16]. However, in case of drift by change in enzyme order, an additional theoretical step is necessary to formally infer the structure of new intermediary metabolites and new enzymatic reactions before experimental validation. For such approaches, most of the time, the new reaction has never been observed in any organism, so that an approach purely based on a search in a database of known reactions is doomed to failure [17]. It is necessary to introduce a knowledge-based approach that implements reasoning in the manner of a biochemist. Such strategies have already been used for designing experimental setup for the analysis of auxotrophic mutants in yeast [18] or for synthetic biology [19]. To be successful, approaches of scientific discovery based on artificial intelligence techniques necessitate close and iterative interactions between chemists, biologists and bioinformaticians [20-22]. This has already led to promising results in the field of drug screening for neglected tropical diseases [23]. To further test the hypothesis of metabolic pathway drift, we decided to combine GSM reconstruction and metabolic profiling, the latter based on a bibliographic survey and mass spectrometry analyses, in an emerging model, the red alga *Chondrus crispus*.

*C*. *crispus* is a red seaweed that has been subject to biological studies for more than two centuries [24]. Its genome was sequenced, and annotation was performed with a focus on metabolic features [25]. A non-exhaustive bibliographic search enabled us to find 15 papers mentioning the identification of metabolites from *C*. *crispus* by various methods of chemical profiling. Nine of them were specifically focused on *C*. *crispus* [26-34], whereas the six others were comparative studies between several algae [35-40]. We selected these papers as a test case for incorporating the bibliographic knowledge into a GSM. Additionally, we decided to acquire additional experimental data regarding two pathways chosen for their complementary interest: sterols and mycosporine-like amino-acids (MAAs). The sterol pathway is well investigated at the comparative genomics level [41] and consists mainly of oxidoreductions on a known skeleton, the sterane, consisting of three hexacarbon rings on one pentacarbon ring [42]. Analytical standards are available for different molecules, enabling level 1 metabolite identification by mass spectrometry, according to the metabolomics standard initiative [43]. MAA synthesis involves combination of different building blocks, and analytical standards are lacking for this class of compounds, limiting metabolite identification to level 2 in best cases [43]. Here, using a logical representation of molecules and reactions, we propose the development of an analogy reasoning model and the use of a generic solver to produce all possible inferences. Integrating this with a global analysis of the genome-scale metabolic network, with targeted experimental profiling, and with comparative genomic analysis, we are able to propose an exhaustive model for two metabolic pathways in *C*. *crispus*, structurally shaped by metabolic pathway drift.

## Results

### Chemical identification of main sterols in *C*. *crispus*, but not of some plant-like biosynthetic intermediates by targeted GC-MS profiling

Results of targeted profiling of 15 sterols plus one immediate precursor (squalene) are summed up in Table 1.

**Table 1.** List of sterols profiled in this study, and comparisons with previous studies. For each compound, analytical parameters (retention time and m/z ratio) are given in S1 Table.

| Analysed compounds | Chemical formula | Found in this study | Previous evidence |
| --- | --- | --- | --- |
| brassicasterol | C<sub>28</sub>H<sub>46</sub>O | yes | [44] (GC-MS), [29] (TLC, GC-MS) |
| campesterol | C<sub>28</sub>H<sub>48</sub>O | yes | [29] (TLC, GC-MS) |
| cholesterol | C<sub>27</sub>H<sub>46</sub>O | yes | [44] (TLC, GC-MS), [29] (TLC, GC-MS) |
| cycloartanol | C<sub>30</sub>H<sub>52</sub>O | no | not reported |
| cycloartenol | C<sub>30</sub>H<sub>50</sub>O | no | [44] (TLC), [35] (TLC) |
| cycloeucalenol | C<sub>30</sub>H<sub>50</sub>O | no | not reported |
| 7-dehydrocholesterol | C<sub>27</sub>H<sub>44</sub>O | yes | [29] (TLC, GC-MS) |
| desmosterol | C<sub>27</sub>H<sub>44</sub>O | yes | [44] (TLC, GC-MS), [45] (GC-MS) |
| ergosterol | C<sub>28</sub>H<sub>44</sub>O | no | not reported |
| fucosterol | C<sub>29</sub>H<sub>48</sub>O | no | not reported |
| lanosterol | C<sub>30</sub>H<sub>50</sub>O | no | [44] (TLC) |
| lathosterol | C<sub>27</sub>H<sub>46</sub>O | yes | [45] (GC-MS) |
| β-sitosterol | C29H50O | yes | [44] (GC-MS), [29] (TLC, GC-MS) |
| squalene | C30H50O | yes | not reported |
| stigmasterol | C29H48O | yes | [44] (GC-MS), [29] (TLC, GC-MS) |
| zymosterol | C27H44O | no | not reported |

In addition to confirm the presence of eight previously identified sterols (brassicasterol, campesterol, cholesterol, 7-dehydrocholesterol, desmosterol, lathosterol, β-sitosterol and stigmasterol), we identified here for the first time one immediate precursor, squalene (S1 Fig) i n *C*. *crispus*. However, we did not find evidence for some other putative intermediates (cycloartanol, cycloeualcenol, ergosterol, fucosterol and zymosterol) that may have been present based on the knowledge of sterol synthesis pathway in other eucaryotes [41, 46]. We also did not find cycloartenol in *C*. *crispus* extracts despite the fact that we are able to identify the cycloartenol standard when added in algal extract (S2 Fig). Cycloartenol has been reported a long time ago in *C*. *crispus* extracts from Roscoff using another analytical technique, thin-layer chromatography (TLC) [35]. However, the same group was unable to isolate cycloartenol from another red alga, *Rytiphlea tinctoria*, using GC-MS [47]. Independently, Saito and Idler [44] isolated lanosterol instead of cycloartenol from *C*. *crispus* using TLC, but also failed to find lanosterol using GC-MS. More recently, a cycloartenol synthase from the red alga *Laurencia dendroidea* was cloned and expressed in yeast cells, where it is able to transform squalene into cycloartenol, but the authors did not report cycloartenol identification in the whole alga by GC-MS, as they did for cholesterol (Calegario et al., 2016). Even if undetectable using GC-MS, another indirect argument for cycloartenol as a biosynthetic intermediate is the presence of a compound with a cyclopropyl ring in another florideophyte red alga, *Tricleocarpa fragilis* [48]. The cyclopropyl ring on sterols is usually made by oxidosqualene cyclisation, and the only described product of this reaction is cycloartenol, so we consider more parsimonious to hypothesize that cycloartenol is below the detection limit rather than considering that this step is performed by an unknown intermediate.

### An unknown compound among most abundant MAAs in *C*. *crispus*

Results of LC-MS targeted profiling of mycosporin-like aminoacids are summed up in Table 2.

Using LC-MS profiling, we confirmed, consistently with previous studies (see references in Table 2), the presence of six mycosporine-like aminoacids in *C*. *crispus*: asterina-330, palythene, palythine, palythinol, porphyra-334 and shinorine. Additionally, we identified mycosporine-glycine for the first time in *C*. *crispus*, and also found a peak at m/z=271.1 that does not match with any already identified candidate MAA, that we named it MAA1 in Table 2. We also decided not to assign the peak at m/z=302,3117 to palytinol, as done previously [49, 51], based on logical reasoning about this part of the pathway (see below). That is the reason why an other unknown compound, MAA2, appears in the table. The relative abundance of MAAs seems to vary according to the sampling dates (Fig 2).

**Table 2.**
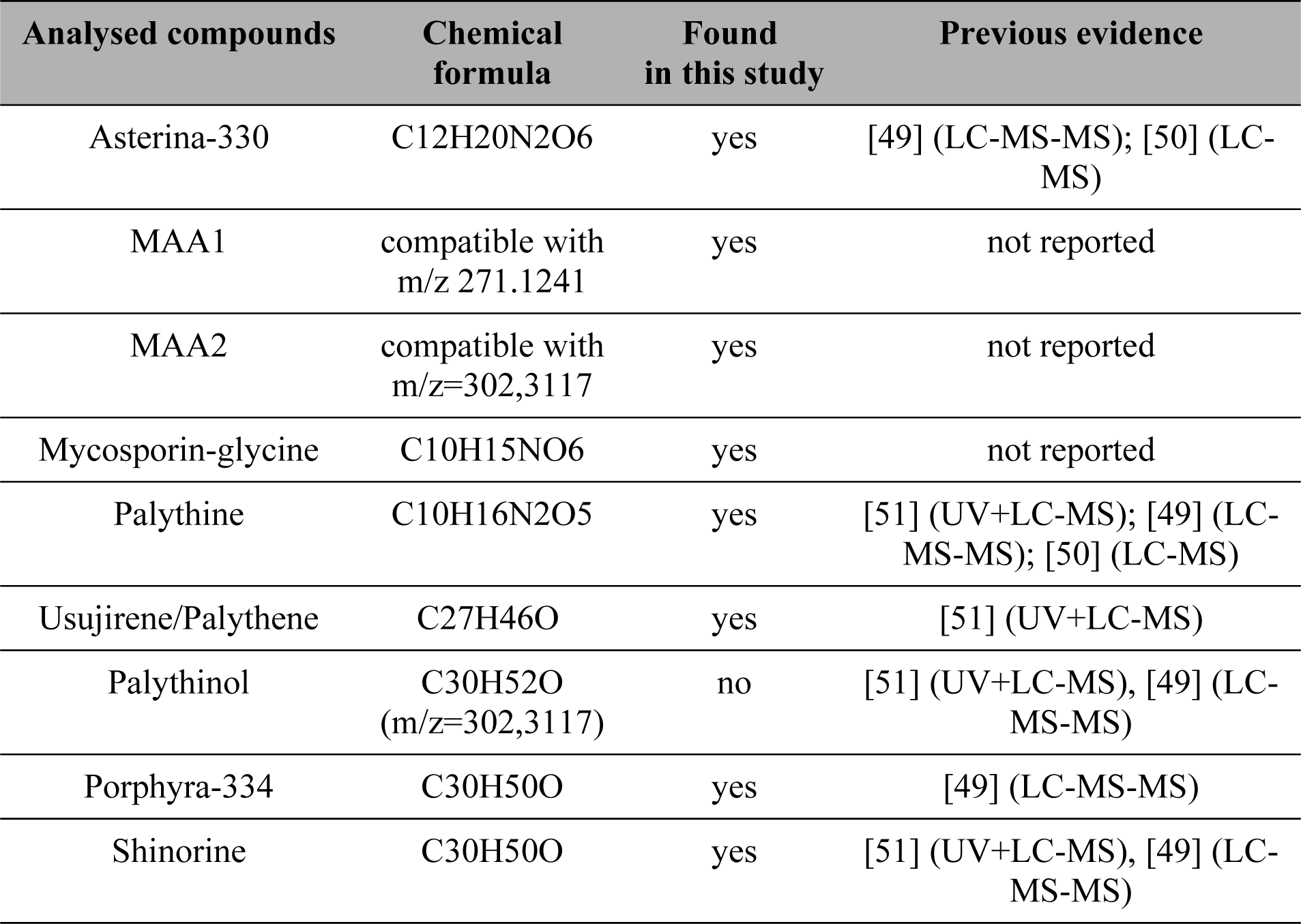
List of mycosporin-like amino-acids identified in this study, and comparisons with previous ones. For each compound, analytical parameters (RT, mz and UV absorption parameters) are given in S2 Table.

**Fig 2.**
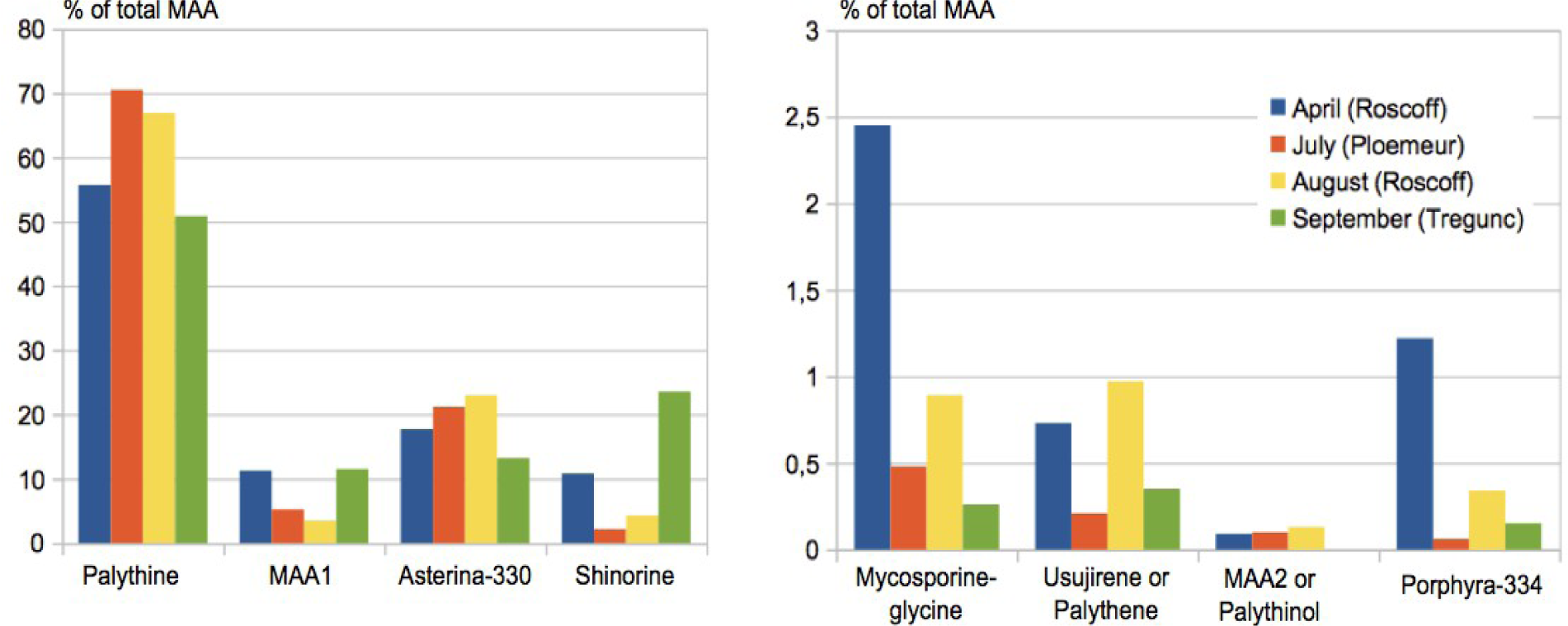
Composition and seasonal variation *(MS quantification)* of MAAs in *C*. *crispus*.

These results should be interpreted carefully because MAAs are known to react differently to MS ionization. Furthermore, under UV, the molar extinction coefficients are different. This allows only semi-quantitative measurements. However, our results are consistent with an independent report of MAA variation in the Galway Bay, Ireland [50]. In both cases, palythine was the most abundant compound. Depending on localisation, and time, then shinorine and asterina-330 were the most abundant compounds, and porphyra-334 was very scarce. The unknown compound at m/z 271.1241, which was here labelled MAA1, is the fourth most abundant MAA in Brittany samples from *C*. *crispus*.

Our new metabolite profiling data on sterols and MAAs were pooled with results from other studies, retrieved by bibliographic search, in order to obtain a set of metabolite targets that was used to constrain the genome-scale reconstruction of the *C*. *crispus* metabolic network (S3-S4 Tables).

### Several non-orthologous genes encode best candidate enzymes for performing conserved reactions in the sterol synthesis pathway

In order to enable comparisons with the automated genome-scale reconstruction, and to facilitate integration with metabolic profiling data, we carried out a comparative genomic analysis of the enzymes involved in the sterol synthesis pathways. Results are summed up in Table 3.

**Table 3.** Comparative genomic analysis of sterol synthesis enzymes. In the first column, the color code for enzymatic steps follows Desmond and Gribaldo [41]. In the four other column, dark blue indicates orthologous sequences, light blue indicates paralogous ones, and green indicates yeast enzymes non orthologous to animal or plant sequences but known to perform the same enzymatic reaction. Five corrected sequences and new predictions are indicated with an asterisk (*) and provided in S1 Dataset.

| Steps | Yeast | Human | *Arabidopsis* | *C. crispus* |
| --- | --- | --- | --- | --- |
| squalene monooxygenation | ERG1 | SQLE | SQE1-7 | scaffolds 90*, 20*, 57* |
| oxydosqualene cyclisation | ERG7 | LSS | CAS | CHC_T00008265001 |
| C-14 demethylation | ERG11 | CYP51A1 | CYP51G1 | CYP51G1 (CHC_T00009303001) |
| C-14 reduction | ERG24 | TM7SF2 | FK | CHC_T00003466001 |
| C-4 demethylation | ERG25 | SC4MOL | SMO1, SMO2 | CHC_T00010320001, scaffold212* |
| delta-8, delta-7 isomerisation | ERG2 | EBP | HYD1 | CHC_T00001257001 |
| C-5 desaturation | ERG3 | SC5DL | STE1 | CHC_T00006481001 |
| C24 or C24' methylation | ERG6 | - | SMT1, SMT2 | CHC_T00009101001, CHC_T00000837001 |
| delta-7 reduction | - | DHCR7 | DWF5 | CHC_T00006492-3001* |
| delta-24 reduction | ERG4 | DHCR24 | DWF1/SSR | CHC_T00002789001 |
| C-22 desaturation | ERG5 | - | CYP710 | CYP805A1-C1 or CYP808A1-H1 |
| cyclopropylsterol isomerisation | - | - | CPI1 | CHC_T00002985001 |

In line with previous analyses on these gene families in eukaryotes [41] or more specifically in green plants [46], the candidate sterol synthesis enzyme set shows a mixture of conservation and divergence. Seven enzymes are encoded by genes that are conserved as 1:1 orthologs, whereas four of them either underwent lineage-specific duplications (squalene epoxidase and C-4 demethylase) or were lost and may have been replaced by distant paralogs (C24 and C24’ methylases and C22 desaturases). In one case, we found no homolog of known plant or animal enzymes performing delta-7/delta-8 isomerisation at all in the *C*. *crispus* genome, but we found a 1:1 ortholog of ERG2, the gene that secondarily took up this function in yeast [41]. We consider this gene the best candidate to test among known gene families, but it is also possible that this reaction is performed by an enzyme encoded by a taxonomically-restricted orphan gene, which have been shown to have some biological roles in other lineages [52]. Actually, this is likely the case in the sterol synthesis pathway in some diatoms, where the epoxisqualene cyclase, otherwise conserved in eucaryotes, was secondarily lost and replaced by another yet unidentified enzyme [53].

We did not carry out a similar genomic analysis for MAAs, because it was already fully done during the annotation of the *C*. *crispus* genome [25], and was recently put in a comparative perspective following the annotation of the MAA genes in an other red alga, *Porphyra umbilicalis* [54].

### Integration of genome-scale reconstruction and targeted chemical profiling highlights the need for *ab initio* inferences to fill knowledge gaps

A global overview of the procedure used to build an integrated metabolic network model for *C*. *crispus* is shown in Fig 3. The network is browsable at: *http://gem-aureme.irisa.fr/ccrgem/index.php/Main_Page*

**Fig 3.**
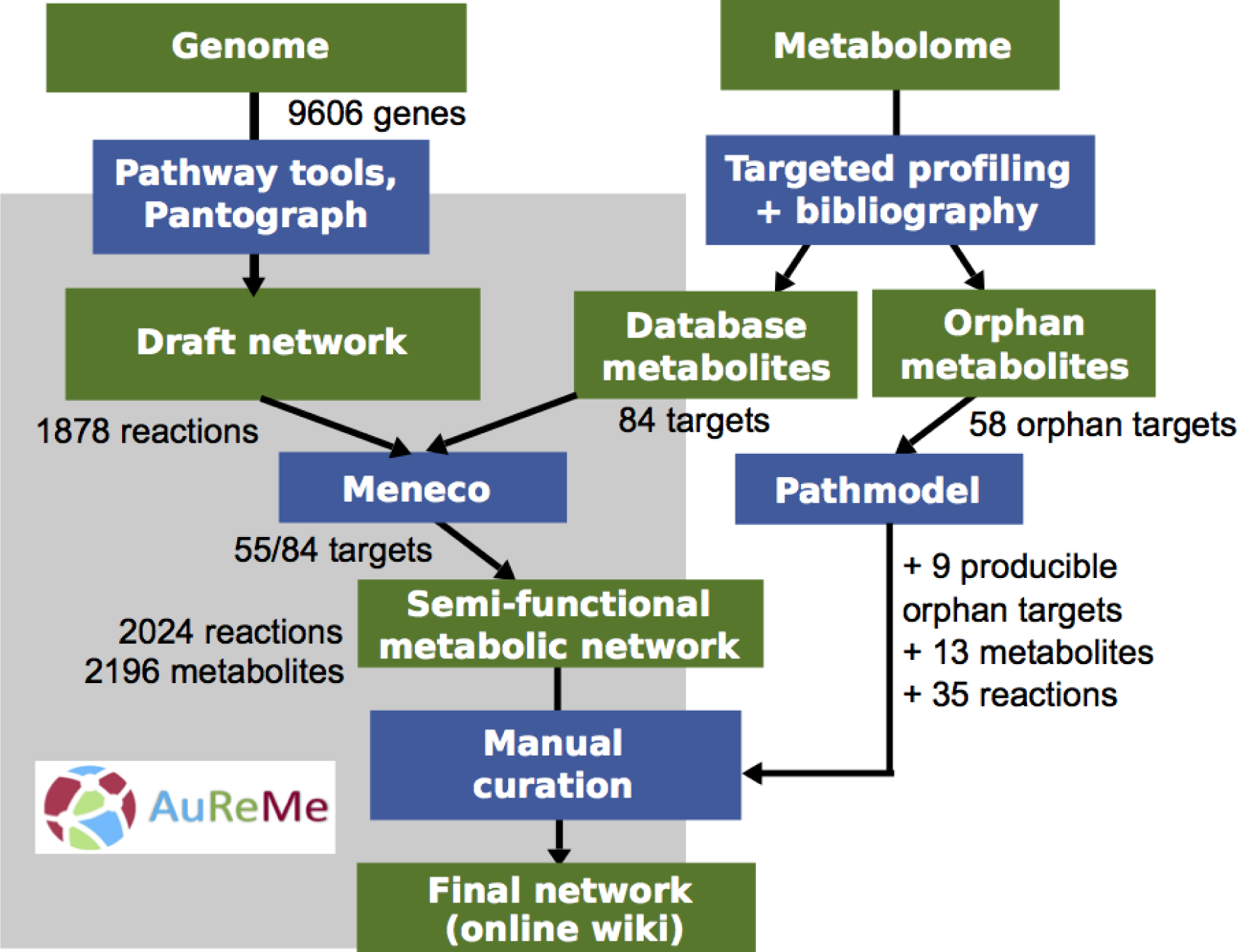
Reconstruction scheme for the genome-scale metabolic network of *C*. *crispus*. Green boxes indicate starting data and resulting new knowledge. Blue boxes indicate the tools that were used to analyse and integrate genome and metabolome data. The part overshadowed in grey indicates tools that are already integrated in the AuReMe workflow [55].

The final network contains 595 reactions coming directly from genome annotation through PathwayTools, 383 reactions coming from orthology with *Arabidopsis thaliana*, 1361 reactions coming from orthology with *Galdieria sulphuraria*, and 1161 reactions coming from orthology with *Ectocarpus siliculosus*. The total number of reactions in the fused network (2024) is in the same range as in the networks of two other macroalgae, *E*. *siliculosus* (1977) and *E*. *subulatus* (2074), reconstructed also using the AuReMe toolbox (Table 4).

**Table 4.** Comparison of global features of genome-scale metabolic networks from macroalgae and other chlorophyllian eucaryotes.

| Species | Reactions | Enzymes | Metabolites | Pathways | Reference |
| --- | --- | --- | --- | --- | --- |
| *C. crispus* | 2024 | 2006 | 2196 | 1108 | This study |
| *E. siliculosus* | 1977 | 2281 | 2132 | 1101 | [55] |
| *E. subulatus* | 2074 | 2445 | 2173 | 1083 | [56] |
| *A. thaliana* | 1567 | 1419 | 1748 | 796 | [57] |
| *C. reinhardtii* | 3083 | 1355 | 1133 | 522 | [58] |

Detailed manual comparisons of the networks from *E*. *siliculosus* and *E*. *subulatus* have shown that all the differences between them are due to technical biases during the reconstruction process [56]. The *C*. *crispus* network once again illustrates the high sensitivity of the results to the quality of the input data. Due to differences in the annotation level, annotation-based reconstruction gave different results between the two *Ectocarpus* species (1661 or 1779 predicted reactions) and in *C*. *crispus* (595 predicted reactions), while orthology-based transfer of central metabolism reactions from *Arabidopsis thaliana* led to a similar number of reactions in all three algae (383 in *C*. *crispus*, 440 in *E*. *siliculosus* and 421 in *E*. *subulatus*). More than half of the reactions (1361 out of 2024) were transferred based on orthology from the red microalga *Galdiera sulphuraria*, which was selected after inspection of the automatically reconstructed annotation-based network available in the MetaCyc database. This illustrates the usefulness of the AuReMe pipeline to efficiently correct for annotation biases using orthology information. Interestingly, orthology-based transfer of reactions from *E*. *siliculosus* to *C*. *crispus* was more successful than orthology transfer from *A*. *thaliana* (1161 versus 383 reactions), despite the fact that *A*. *thaliana* is more closely related to *C*. *crispus* than to *E*. *siliculosus*. This clearly shows how technical issues interfere with biology: the higher number of transferred reactions from *E*. *siliculosus* is linked with the fact that, since both networks being already incorporated in the AuReMe pipeline through the PADMet format, correspondences of reactions IDs were easier than with the *Arabidopsis* network which was reconstructed using a different workflow [57]. Variability comes also from the level of database completeness: the increase of database completeness with time is striking when comparing the reaction numbers between *A*. *thaliana* [57] and *C*. *reinhardtii* [58].

Another important comparison level is the number of metabolites for which the presence in *C*. *crispus* is experimentally proven. 84 metabolites from *C*. *crispus* were indexed in the MetaCyc database and could thus be used as targets, that should be present in the final network. Using the gap-filling program Meneco, we managed to incorporate 55 of them, which is higher than the two *Ectocarpus* species (50 targets), but still only represents two thirds of all targets. In addition, there were 58 orphan metabolites that could not be incorporated automatically, because they were not yet indexed in MetaCyc. This prompted us to develop Pathmodel, a new tool that enables to infer new metabolic reactions and new molecules to connect orphan metabolites with the main network. This tool was tested on the sterol and mycosporine-like amino-acid synthesis pathways because they were suitable to address complementary issues. The sterol pathway raised the problem of connecting and integrating various portions of known sterol synthesis pathways from animals and plants (Fig 4A) while the MAA pathway raised the problem of integrating unannotated compounds that were identified uniquely based on their m/z ratio (Fig 4B).

**Fig 4.**
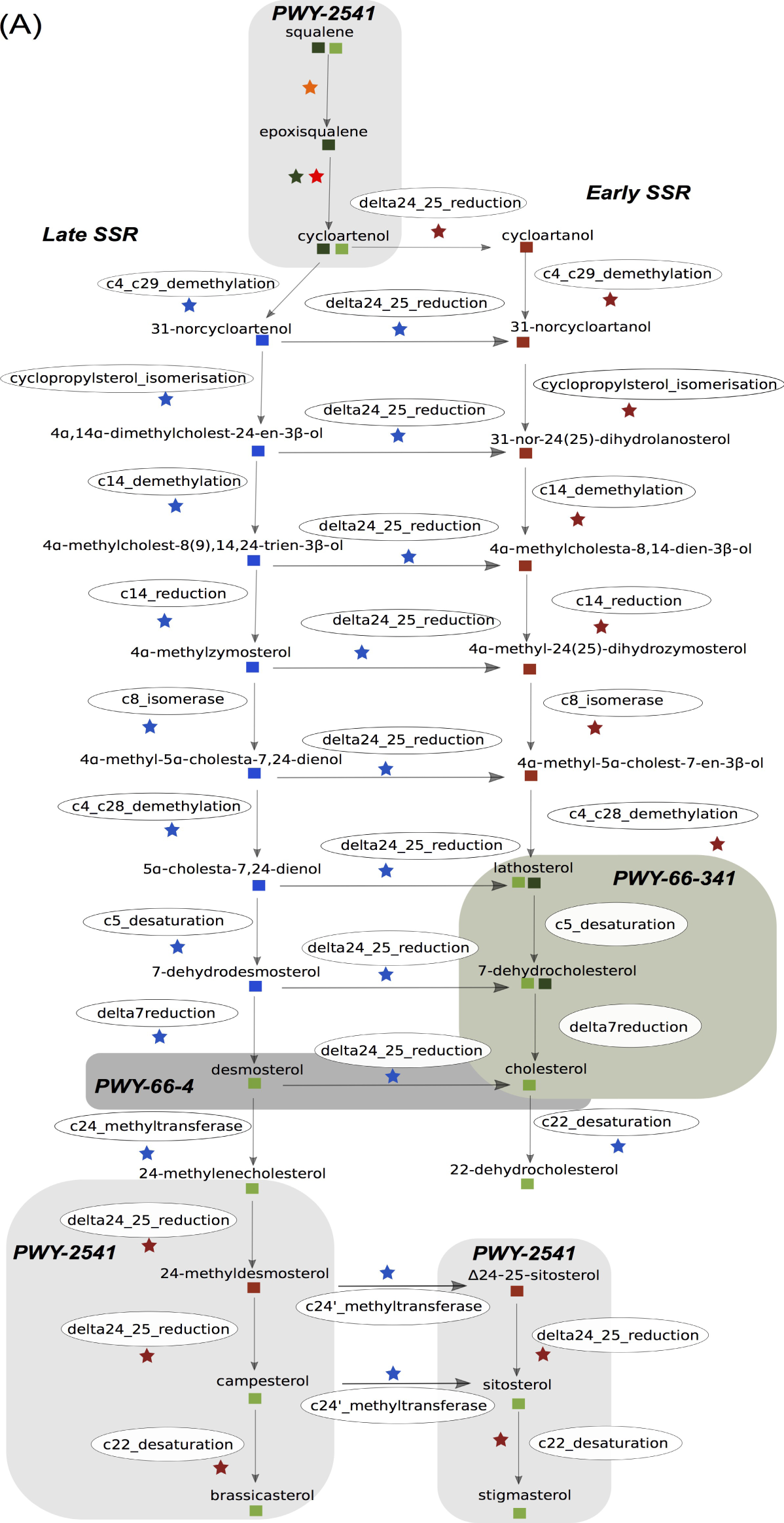
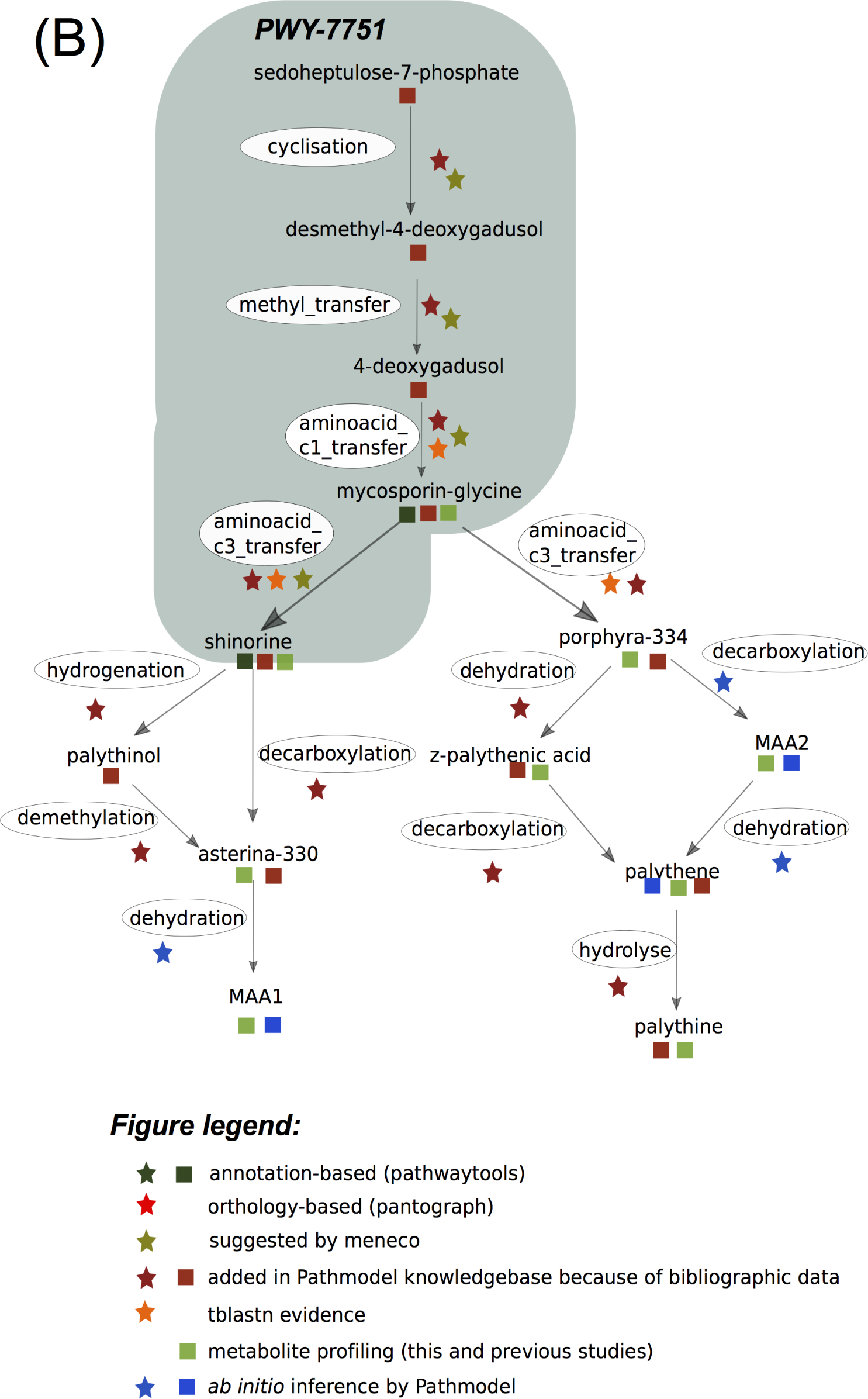
Overview of pathways reconstructed with Pathmodel using multiple heterogenous data and analogical reasoning. (A) Sterol biosynthesis pathway. Portions that are already in the MetaCyc database are highlighted with grey boxes. PWY-2541: plant sterol biosynthesis pathway. PWY-66-34: animal modified Kandutsch-Russell pathway. PWY-66-4: animal Bloch pathway. See part (B) fore the detailed figure legend. (B) MAA Biosyntesis pathway. PWY-7751: shinorine biosynthesis pathway. The figure legend details the various data sources integrated to infer the pathways. Stars indicate reactions, squares indicate molecules.

The metabolites present in *C*. *crispus* only partially fitted with standard pathways indexed in the MetaCyc database, for different reasons. Regarding the sterol synthesis pathways, they belong to three different pathways: cycloartenol, 24-epicampesterol, brassicasterol, sitosterol and stigmasterol belong to the canonical plant sterol biosynthesis pathway (PWY-2541; [59]), whereas lathosterol, 7-dehydrocholesterol belong to the animal modified Kandutsch-Russell pathway (PWY-66-341, [60]) and desmosterol belongs to the animal Bloch pathway (PWY-66-4, [60]). Both animal pathways result in the production of cholesterol at their end. Additionally, 22-dehydrocholesterol is not a part of any of those pathways. Regarding MAAs, mycosporin-glycin and shinorine belong to the shinorine biosynthesis pathway (PWY-7751), corresponding to the best understood part of the pathway [61], but all other compounds identified in *C*. *crispus* are absent from the MetaCyc database. Morover, the query of public chemical structure databases cannot help in assigning a tentative structure to the peak corresponding to MAA1. All those limitations explain why we selected those two pathways as case studies to develop the Pathmodel method to infer *ab initio* new reactions and new metabolites.

### Inferring new metabolic reactions using Answer Set Programming

The Pathmodel method takes as input a knowledge base including a set of known metabolites, a set of observed mass-to-charge (m/z) ratios for unknown metabolites, and a set of known enzymatic reactions. For each pair of metabolites which are not linked by a reaction in the knowledge base, the method checks whether a type of known reaction can occur between them, and further derives from known reactions new candidate metabolites corresponding to observed unassigned mass-to-charge ratios. This is the basis for the selection of new reaction occurrences and/or new metabolites, using either deductive or analogical reasoning (Fig 5).

**Fig 5.**
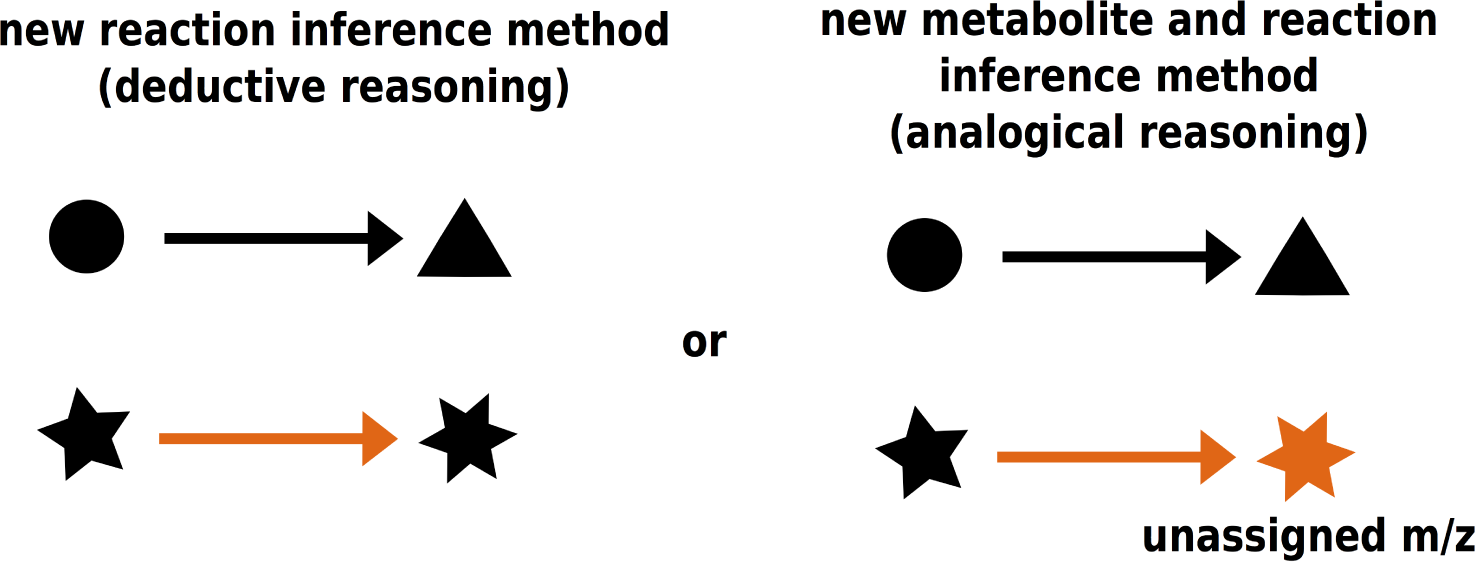
The two reasoning methods implemented in Pathmodel. Input data encoded in the knowledge base are in black, newly inferred reactions and metabolites structures are in orange.

Molecules are modeled by a set of logical predicates *atoms* (identified by a number and atom types) and *bonds* (identified by atom numbers and bond type), as highlighted in orange on Fig 6.

**Fig 6.**
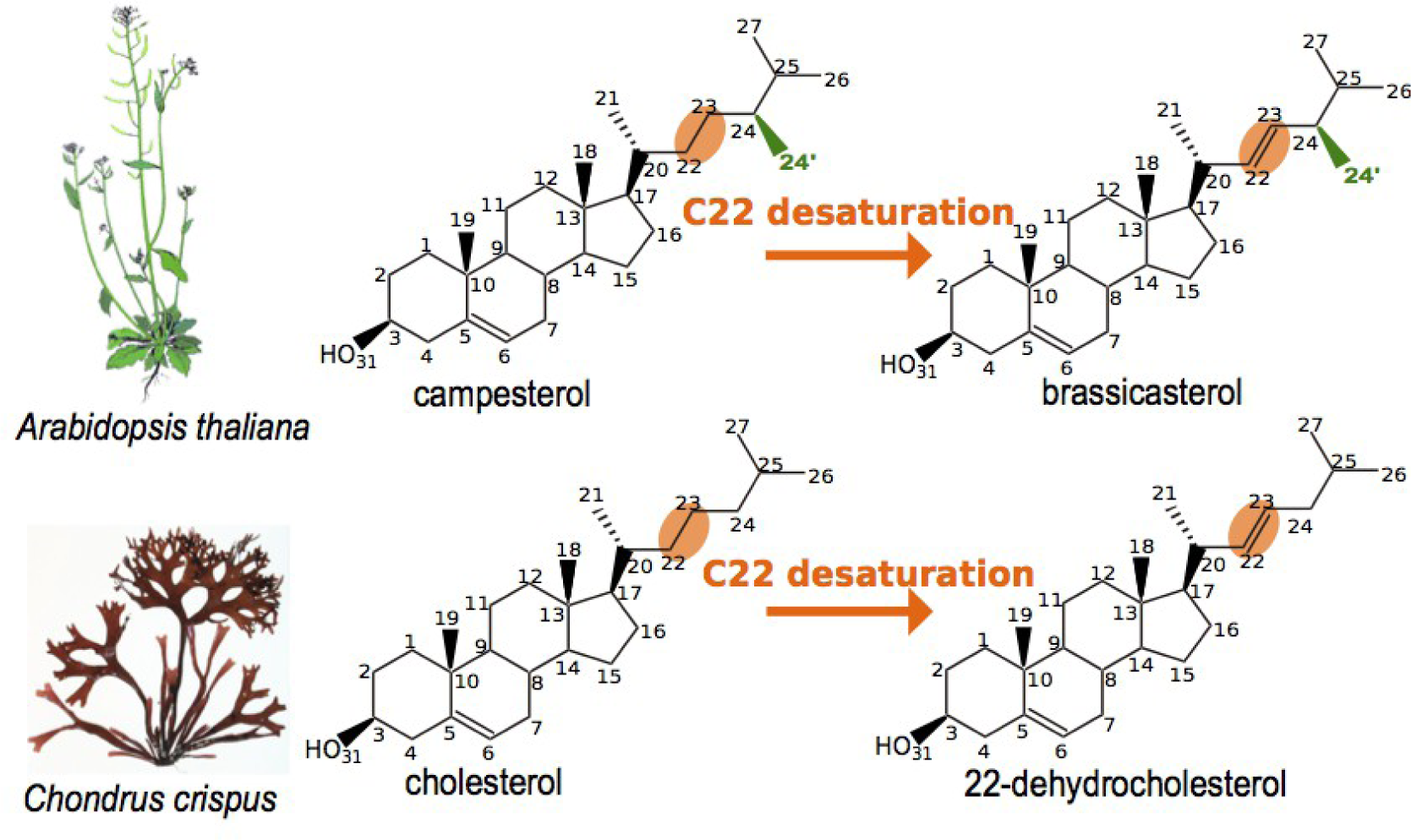
Detailed encoding of metabolites and reactions in Pathmodel. In black, molecules structures with carbon and oxygen atoms labelled. For example, carbon 22 from brassicasterol is encoded by the predicate *atom(“brassicasterol”,22, carb)*. In orange, position of the bond between atoms submitted to the chemical reaction, encoded by the predicate *bond (“brassicasterol”, double,22,23)*. In green: the C24 methyl group that makes the difference between molecules from *Arabidopsis thaliana* and from *Chondrus crispus*.

In order to perform this reasoning, the program needs some preprocessing steps (S4 Fig). For each newly inferred molecule, the theoretical m/z ratio is determined by logical rules, which was encoded in the program MZComputation.lp. First, the number of hydrogens for each atom of a molecule was deduced (predicate *numberHydrogens)* from the total number of bonds in which the atom is involved and from the valence of the atom. Then the number of each atom species (hydrogens, carbons,…) is determined for each molecule (predicate *moleculeComposition*). Finally, the m/z ratios are derived from the molecular composition (predicate *moleculeMZ*), using the following formula:

moleculeMZ (MoleculeName, MassCarbon*NumberCarbon + MassHydrogen*NumberHydrogen + MassOxygen*NumberOxygen + MassNitrogen*NumberNitrogen + MassPhosphorus*NumberPhosphorus):-moleculeComposition(MoleculeName, NumberCarbon, NumberHydrogen, NumberOxygen, NumberNitrogen, NumberPhosphorus).

In this formula, atomic weights are encoded following the latest IUPAC Technical Report [62], truncated after the fourth decimal and multiplied by 1000 because the ASP syntax does not allow the use of decimals.

The predicate *reaction* models the link between two molecules (a reactant and a product, e.g. *reaction(c22_desaturation,“24-epicampesterol”, “brassicasterol”)*). By comparing reactants and products, the program ReactionSiteExtraction.lp characterizes two structures of the reaction site containing atoms and bonds involved in the reaction: one structure describes the reaction site before the reaction (Figure 6, simple bond between atoms 22 and 23 in campesterol) and the other describes the reaction site after the reaction (Fig 6, double bond 22-23 in brassicasterol). Predicates *diffAtomBeforeReaction, diffBondBeforeReaction, diffAtomAfterReaction and diffBondAfterReaction* compare atoms and bonds between the reactant and the product and extract the two structures. Then these two structures are compared to the structure of all other molecules in the knowledge base (predicates *siteBeforeReaction* and *siteAfterReaction)*. These predicates characterize sub-structures of the molecules that can be part of a reaction. These are the bases for the selection of new potential reactants or products and the inference by a reasoning component of new reaction occurrences or new metabolites, using either deductive or analogical reasoning in the PathModel.lp program.

By deductive reasoning, the reference molecule pair of each reaction (Fig 6, campesterol and brassicasterol) is compared to the structures of a potential reactant-product pair sharing a common chemical structure (Fig 6, cholesterol and 22-dehydrocholesterol, sharing a sterane skeleton) with the predicate *deductiveReasoningInference*. The presence of the reaction site in the two putative molecules is checked by using the predicates *siteBeforeReaction* and *siteAfterReaction*. Furthermore, if the product and the reactant have the same overall structure, except for the reaction site (see Fig 6, bond between atoms 22 and 23), the program will infer that the reaction actually occurs between the reactant and the product (Fig 6, desaturation between cholesterol and 22-dehydrocholesterol). To constraint further the number of possible pathways, a predicate *absentmolecules* was added to avoid pathways going through compounds for which targeted profiling with analytical standards gives strong evidence for real absence (here ergosterol, fucosterol and zymosterol).

By analogical reasoning, all possible reactions are applied to potential reactants, and resulting products are filtered using their structures and m/z ratios. The predicate *newMetaboliteName* creates all the possible products from a known molecule using all the reactions in the knowledge base. These possible metabolites are filtered using their m/z ratios, which must correspond to an observed m/z ratio (predicate *possibleMetabolite*) and checked if they share the same structure as a known molecule (predicate *alreadyKnownMolecule*).

Given a source molecule and a target molecule, the program will take several inference steps iteratively applying either analogical or deductive reasoning modes. To connect the source and the target molecules along a pathway, Pathmodel infers missing reactions and metabolites using a minimal number of reactions.

## Discussion

### Multiple alternative pathways for sterol synthesis

Based on the available genomic and metabolomic data, we can propose two alternative pathways from cycloartenol to cholesterol, depending on when the side-chain reductase (SSR) enzyme is acting (Fig 7).

**Fig 7.**
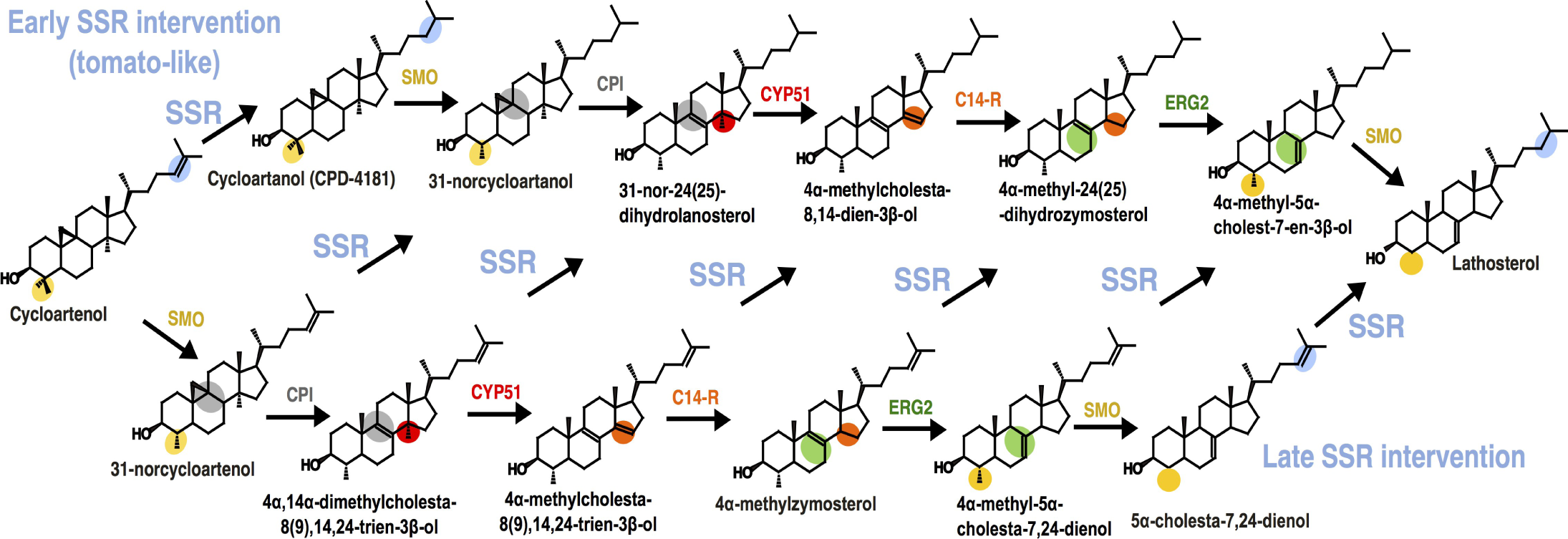
Alternative lathosterol synthesis pathways from cycloartenol in *C*. *crispus*. Enzyme names refer to terrestrial plants, escept for ERG2 that refers to yeast, and are explained in Table 3.

The « early SSR » pathway is based on the model previously published for tomato [46]. The reactions were manually incorporated in the PathModel knowledge base because they are not yet available in MetaCyc. If *C*. *crispus* uses this pathway, the metabolic intermediates would be identical to tomato, but there would be an important difference concerning the enzymes. Indeed, the genes encoding SSR are duplicated in Solanaceae (tomato and potato) but not in other plants, in the *C*. *crispus* genome or in any red algal genome analyzed so far (Supp Figure 3). In Solanacea, SSR2 acts on cycloartenol whereas SSR1 acts late in the phytosterol synthesis pathway, as does the unduplicated SSR from non-solanaceous plants. Moreover, SSR is known to be catalytically promiscuous, and in humans the unique SSR enzyme is able to act either late or early [60]. Therefore, our data suggest that the single SSR is also flexible in *Chondrus* as it is in humans, enabling the existence of multiple synthesis pathways leading to cholesterol. A high level of reticulation with multiple alternative routes in the plant sterol synthesis pathways has already been suggested [59], although only the main pathway has been incorporated in the knowledge base (see for example PWY-2541 in MetaCyc). Consistently, Pathmodel suggested that SSR could act on all possible intermediates. However, flux analyses in mouse have shown that among all theoretical possibilities, two distinct pathways, whose relative abundance vary accross tissue, are sufficient to enable refined and partially distinct regulations [60].

Another major difference with land plants concerns the position of the sterol methyltransferases, that are necessary to produce methylated sterols like campesterol or brassicasterol. In the standard model for land plants, a first methylation occurs directly on cycloartenol whereas the second one occurs later on 24-methylenelophenol [59]. Although this possibility cannot be fully ruled out concerning *C*. *crispus* with present data, various pieces of evidence points toward the necessity to consider alternative pathways. First, we did not find any evidence for the presence of cycloeucalenol or fucosterol, which are common synthetic intermediates in the plant pathway. Second, another methylated sterol, 24-methylenecholesterol, was identified previously in *C*. *crispus* [29]. In line with this, and building on other reports about methyltransferase catalytic promiscuity accross land plants and green algae, [63, 64], Pathmodel inferred an alternative synthesis pathway for methylated sterols through C24-methylation on desmosterol (Fig 4). This option highly reduces the number of non-identified methylated intermediates, limiting them to 24-methydesmosterol and Δ24-25-sitosterol. It seems also more relevant from a quantitative viewpoint, because this late methylation step would enable the production of methylated sterols using the late SSR pathway, which is also in agreement with the formation of cholesterol as the main sterol.

### New candidate enzymes for decarboxylation and dehydration lead to a more consistent model for MAA synthesis pathway in *C*. *crispus*

The upstream part of the MAA synthesis pathway in *C*. *crispus*, down to shinorine and porphyra-334, follows the current consensus. For this part, candidate enzymes were already proposed [54], and this knowledge is already partially incorporated in the MetaCyc database (PWY-7751 on Fig 4). Here we added further experimental support to the presence of this part of the pathway, performing the first identification of mycosporine-glycine in *C*. *crispus* (Fig 4 and Fig 8). We also encoded in the Pathmodel knowledge base an extended version of the aminoacid C3-transfer reaction (RXN-17371 in MetaCyc) to incorporate the already formulated hypothesis, based on structural comparisons between molecules, that MysD can also perform the aminoacid C-3 transfer of threonine, thus leading to porphyra-334 (Fig 8, reaction in red, redrawn from [54]). For the more downstream part of the pathway, some reactions have been proposed, such as decarboxylation of shinorine to asterina-330, but without association with a specific enzyme family [65]. Encoding this literature-based information and constraining the Pathmodel output to find a pathway leading to a molecule structure compatible with the mesured m/z ratio for MAA1, it was possible to infer the hypothetic structure shown on Fig 8. The reaction leading from asterina-330 to MAA1 would be a dehydration (in purple), the same kind of reaction that is already observed between other MAAs such as porphyra-334 and Z-palythenic acid, a compound not identified in *C*. *crispus* (Fig 4).

**Fig 8.**
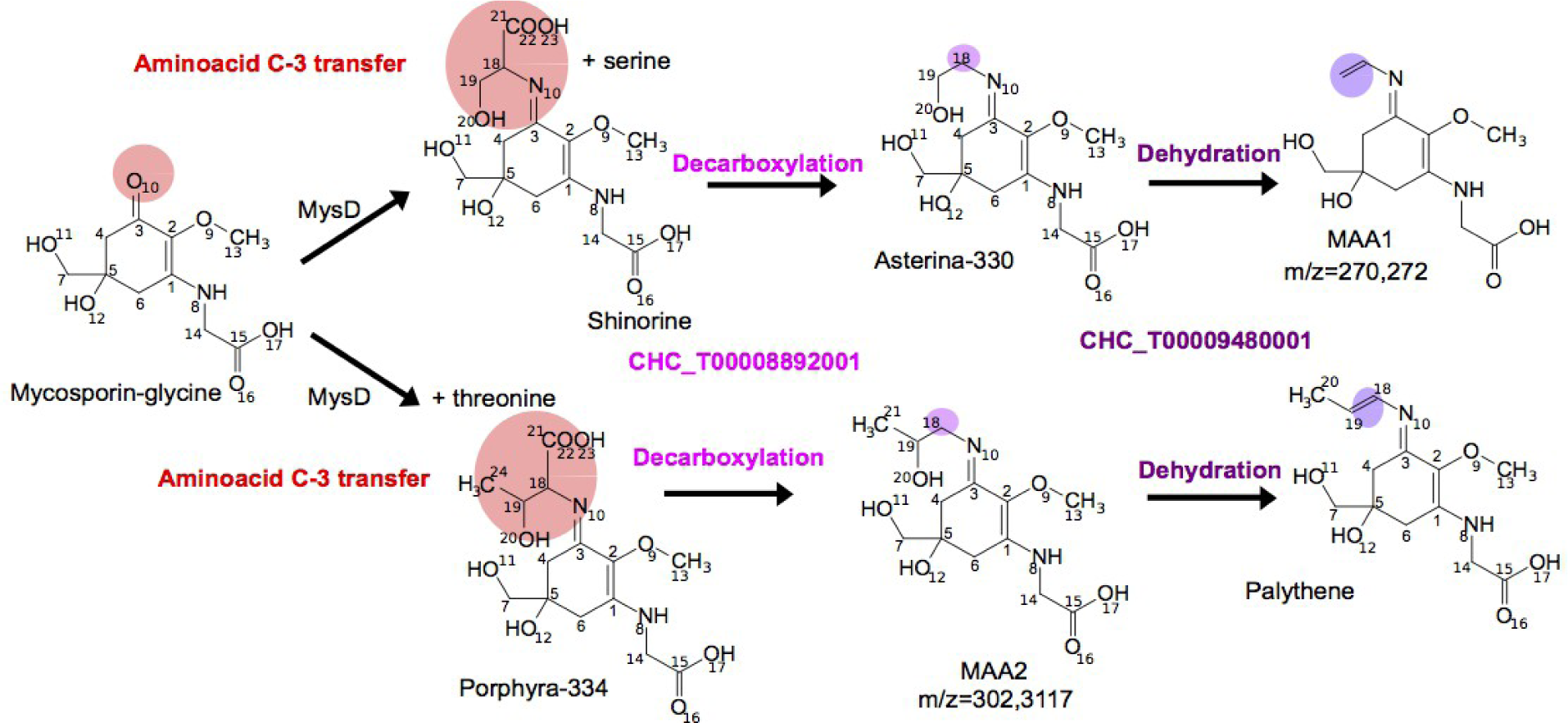
New candidate reactions, enzymes and metabolites in the downstream part of the MAA biosynthesis pathway in *C*. *crispus*. Structure of MAAs and their precursors are drawn with carbon and oxygen atom labelling corresponding to the numeration used in Pathmodel. The two newly inferred reactions are serine/threonine decarboxylation, in pink, and serine/threonine dehydration, in purple.

For both decarboxylation and dehydration reactions, no candidate enzymes were mentioned so far in the literature related to MAA biosynthesis pathways. We thus performed a simple semantic search on a draft version of the GSM from *C*. *crispus*, to identify other enzymes that may perform those reactions on a serine coupled with other chemical building blocks. Serine decarboxylation indeed occurs in phospholipid metabolism and was inferred in the *Chondrus* GSM based on orthology with *Galdieria sulphuraria*. The candidate gene is CHC_T00008892001. Interestingly, there is some evidence of catalytic promiscuity for this enzyme, enabling it to also decarboxylate a threonine residue. So far, biochemical data in mammalian cell cultures indicate that phosphatidylthreonine decarboxylation by phosphatidylserine occurs, but with a weak activity [66]. The *in vivo* occurrence and biosynthetic origin of phosphatidylthreonine were only recently demonstrated using HPLC-MS/MS in the apicomplexan parasite *Toxoplasma gondii* where it is produced by a phosphatidylthreonine synthase coming from an ancient gene duplication of a phosphatidylserine synthase specifically in the lineage encompassing stramenopiles, alveolates and rhizarians [67]. Therefore, we hypothesize that the enzyme may be promiscuous in *C*. *crispus* and may also perform serine/threonine decarboxylation on a serine/threonine linked to a mycosporin-glycin instead or in addition to performing this on a phospholipid.

Following the same rationale, we performed a semantic search for a serine/threonine dehydratase, and found the enzyme encoded by CHC_T00009480001. This enzyme was predicted based on Pathway Tools to be involved either in degradation of glycine betaine, purine nucleobases, or L-serine, and is a member of the Pyridoxal-phosphate dependent enzyme family, that contains both the human serine dehydratase (EC:4.3.1.17; P20132) and the *E*. *coli* threonine dehydratase (EC:4.3.1.19; P04968) signatures. Common ancestry for serine and threonine dehydratases has been proposed long ago [68]. However, it should be noted that enzymatic promiscuity is very high among pyridoxal-phosphate dependent enzymes [69] and that hydrolases represent only a minor fraction of their overall described biochemical activities [70].

Inferring a single pair of enzymes to decarboxylate shinorine and porphyra-334 and further dehydrate their derivatives was also parsimonious in respect to the absence of a peak corresponding to Z-palythenic acid in *C*. *crispus* extracts, which does not support dehydration occuring before decarboxylation, as proposed in other species (Fig 4 and [65]). The structure of a new intermediate was therefore inferred manually, leading to MAA2 on Figure 8. Calculating its m/z ratio, we found this was identical to palythinol, a compound previously considered to be present in *C*. *crispus* based on UV+LC-MS or LC-MS/MS data [49, 51]. We then verified using Pathmodel that constraining the pathway search with a molecule having a m/z ratio of 302,3177 leads to the same actual MAA2 as a proposed unique solution. Because there is no synthesis-based analytical standard available for palythinol, as for all other MAAs, it was useful to make this alternative hypothesis explicit. From a genomic viewpoint, switching palythinol with MAA2 does not necessitate a candidate enzyme to perform hydrogenation and demethylation on a MAA-like substrate (Fig. 4), and thus reduces the number of unassigned enzymatic activities to candidate genes.

### Implications of possible sterol and MAA synthesis pathways in *C*. *crispus* on evolutionary scenarios regarding metabolic pathway drift

Our study demonstrates that data on metabolite occurrence can be explicitly incorporated into the quality criteria for evaluating a GSM. Putting more emphasis on metabolites, especially the missing ones, creates new methodological challenges regarding *ab initio* inferences of pathways when enzymes are not yet known, and we have shown that it is now possible to build new tools to specifically address those challenges. The next issue is about the scalability of our approach. The Pathmodel version we present here is a working prototype that can already be applied to other metabolic pathways in *C*. *crispus* or in other organisms where genomic and metabolomic data are available. Further improvements should be done in order to minimize the user’s burden in manually entering molecular structures. It is not yet possible to fully automate the atom numbering during metabolic reaction. Note that the five main existing solutions have a success rate of 91% compared with manual mapping, which means that errors would remain with such an approach [71]. We have thus proposed a graphical output in order to facilitate the check of encoded molecule structures (S5-S6 Figs).

The Pathmodel tool was developed to support reasoning based on the metabolic pathway drift hypothesis in order to automatically infer new reactions and metabolites. A first key feature of the successful application of this strategy was the precision and the quality of the biochemical and biological knowledge encoded in Pathmodel. Generalizing this approach to any other application will similarly require interactions between chemists, biologists and computer scientists. The second key feature of Pathmodel is to be focused on a selected pathway rather than on a complete genome-scale metabolic network. The selection of the relevant pathway to be considered - for instance from preliminary evidences extracted from metabolomics analysis - is therefore a key pre-processing step to combine and filter the predictions of Pathmodel with genomics and metabolomics data.

Whatever the actual topology of the sterol and MAA pathways in *C*. *crispus*, each discussed hypotheses have implications regarding metabolic pathway drift. All possible sterol pathways provide further strong candidates case studies for a drift by non-homologous enzyme replacement, and the new pathways inferred by Pathmodel provide candidates case studies for of drift by enzyme inversion. The unresolved point with the sterol pathways is that, among eukaryotes, there is no consensus yet about the ancestral order of enzymatic reactions. Experimental data are too disparate across the tree of life to enable firm conclusions on this. In that respect, the MAA pathway is interesting, because if our hypothesis about decarboxylation of porphyra-334 before dehydration is true, this would mean that an enzymatic inversion took place in other lineages where porphyra-334 is first dehydrated to Z-palytenic acid and then decarboxylated to palythene. Here the limit is that, to date, enzymes are unknown for both reactions, so the system is not yet genomically tractable. Identifying close enzymatic inversions is important, because experimental analyses on *E*. *coli* have shown that drastic pathway rewiring by enzyme knockout or gene overexpression can led to toxic intermediates [72]. Enzyme inversion would provide a milder mechanism for gradual divergence of pathways. But to identify such cases we need genomic and metabolomic data for more closely related model species. Such data will become available in the coming years thanks to ongoing integrative sequencing and metabolomic projects.

## Material and Methods

### Sampling of algae

For sterol analyses, samples from *Chondrus crispus* were collected from a population on the shore at Roscoff, France, in front of the Station Biologique (48°43’38’’ N; 3°59’04’’ W). Algal cultures were maintained in 10 L flasks in a culture room at 14°C using filtered seawater and aerated with filtered (0.22 µm) compressed air to avoid CO2 depletion. Photosynthetically active radiation (PAR) was provided by Philips daylight fluorescence tubes at a photon flux density of 40 µmol.m^-2^.s^-1^ for 10 h.d^-1^. The algal samples were freeze dried, ground to powder using a cryogrinder and stored at −80°C.

For MAAs analysis, more than 50 g (wet weight) of *Chondrus crispus* were collected along the Brittany coasts (France) at Ploemeur (47°42’07’’ N; 3°24’31’’ W) in July 2013, Roscoff (48°43’38’’ N; 3°59’04’’ W) in April and August 2013, and Tregunc (47°50’25’’N; 3°54’08’’ W) in September 2013.

### Standards and reagents

Cholesterol, stigmasterol, *β*-sitosterol, 7-dehydrocholesterol, lathosterol (5*α*-cholest-7-en-3*β*-ol), squalene, campesterol, brassicasterol, desmosterol, lanosterol, fucosterol, cycloartenol, 5*α*-cholestane (internal standard) were acquired from Sigma-Aldrich (Saint-Quentin-Fallavier, France), cycloartanol and cycloeucalenol from Chemfaces (Wuhan, China) and zymosterol from Avanti Polar Lipids (Alabaster, USA). The C7-C40 Saturated Alkanes Standards were acquired from Supelco (Bellefonte, USA). Reagents used for extraction, saponification, and derivation steps were *n*-hexane, ethyl acetate, acetonitrile, methanol (Carlo ERBA Reagents, Val de Reuil, France), (trimethylsilyl) diazomethane, toluene (Sigma-Aldrich, Saint-Quentin-Fallavier, France) and N,O-bis(trimethylsilyl)trifluoroacetamide with trimethylcholorosilane (BSTFA:TMCS (99:1)) (Supelco, Bellefonte, USA).

### Standard preparation

Stock solutions of cholesterol, stigmasterol, *β*-sitosterol, 7-dehydrocholesterol, lathosterol (5*α*-cholest-7-en-3*β*-ol), squalene, campesterol, brassicasterol, desmosterol, lanosterol, fucosterol, cycloartenol and 5*α*-cholestane were prepared in hexane with a concentration of 5 mg.mL^-1^. Working solutions were made at a concentration of 1 mg.mL^-1^, in hexane, by diluting stock solutions. The C7-C40 Saturated Alkanes Standard stock had a concentration of 1 mg.mL^-1^ and a working solution was made at a concentration of 0.1 mg.mL^-1^. All solutions were stored at −20°C.

### Sample preparation

Dried algal samples (60 mg) were extracted with 2mL ethyl acetate by continuous agitation for 1 hour at 4°C. After 10 min of centrifugation at 4 000 rpm, the solvent was removed, the extracts were saponified in 3 mL of methanolic potassium hydroxide solution (1M) by 1 hour incubation at 90°C. The saponification reaction was stopped by plunging samples into an ice bath for 30 min minimum. The unsaponifiable fraction was extracted with 2 mL of hexane and 1.2 mL of water and centrifuged at 2000 rpm for 5 min. The upper phase was collected, dried under N2, and resuspended with 120 µL of (trimethylsilyl) diazomethane, 50 µL of methanol:toluene (2:1 (v/v)) and 5 µL of 5*α*-cholestane (1 mg.mL^-1^) as internal standard. The mixture was vortexed for 30 seconds, and heated at 37°C for 30 min. After a second evaporation under N2, 50 µL of acetonitrile and 50 µL of BSTFA:TMCS (99:1) were added to the dry residue, vortexed for 30 seconds and heated at 60°C for 30 min. After final evaporation under N2, the extract was resuspended in 100 µL of hexane, transferred into a sample vial and stored at −80°C until the GC-MS analysis.

For MAAs, one gram of dried algae was extracted twice for two hours under continuous shaking with 10 mL of acetone. After 5 min of centrifugation at 3 000 rpm, acetone was discarded and samples were re-extracted twice with 10 mL water/acetone (30/70, v/v) for 24 hours under continuous shaking at 120 rpm. Water/acetone supernatants were pooled, added to one gram of silica and evaporated to dryness by rotary evaporation. Extracts were then purified by silica gel chromatography column with dichloromethane/methanol mixtures and MAAs were eluted with 200 mL of dichloromethane/methanol (15/85, v/v). After rotary evaporation, samples were re-suspended in water/methanol (50/50, v/v) and filtrated using 0.45 µm syringes filter. Solution were adjusted to a final concentration of 1 mg.mL^-1^ and stored at 3°C until LC-MS analysis.

### Sterol analysis by gas chromatography-mass spectrometry

The sterols were analyzed on a 7890 Agilent Technologies gas chromatography coupled with a 5975C Agilent Technologies mass spectrometer (GC-MS). A HP-5MS capillary GC column (30 m x 0.25 mm x 0.25 µm) from J&W Scientific (CA, USA) was used for separation and UHP helium was used as carrier gas at flow rate to 1 mL.min^-1^. The temperature of the injector was 280°C and the detector temperature was 315°C. After injection, the oven temperature was kept at 60°C for 1 min. The temperature was increased from 60°C to 100°C at a rate of 25°C.min^-1^, then to 250°C at a rate of 15°C.min^-1^, then to 315°C at a rate of 3°C.min^-1^ and then held at 315°C for 2 min, resulting in a total run time of 37 min.

Electronic impact mass spectra were measured at 70eV and an ionization temperature of 250°C. The mass spectra scanned from m/z 50 to m/z 500. Peaks were identified based on the comparisons with the retention times and the mass spectra (S1 Table).

### MAA analysis by liquid chromatography-mass spectrometry

High Resolution Mass Spectrometry was carried out on a microTOF-Q II (Bruker Daltonics, Germany) coupled to an Ultimate 3000 LC System (Dionex, Germany). Experiments were performed on a Gemini C6-Phenyl column (250 mm x 4.6 mm x 5 µm) (Phenomenex, Germany). The gradient was as follows: methanol/water (20:80, v/v) with 0.2% acid acetic for two minutes to 100 % methanol with 0.2% acid acetic in 23 minutes. The UV detector was set to 330 nm, flow rate was kept constant at 0.4 mL.min^-1^ and column temperature set at 30°C. MS spectra were recorded in positive ESI mode with a drying gas temperature of 220°C, a nitrogen flow of 12 L.min^-1^, a nebulizer pressure set to 60 psi, and a collision energy of 20 eV. MAAs were identified by HR-MS on the basis of the detection of the pseudo-molecular ion [M+H]^+^ with a *m/z v*alue varying less than ± 0.02 Da compared to the theoretical *m/z* value. In the absence of commercially available standards, relative quantification of MAAs in each sample was estimated by calculating the ratio between the area under the curve of the Extracted Ion Chromatogram (EIC) corresponding to the selected MAAs and the sum of the areas under the curve of the EIC of all MAAs detected in the algal extract. The same procedure was applied to UV detection (S2 Table).

### Genome-scale metabolic network reconstruction

Genome-scale metabolic network reconstruction was performed using the AuReMe pipeline [55]. A set of 89 targets coming from the literature was used as an input and is provided in S3 Table. Orphan metabolites that are experimentally supported but do not have a MetaCyc ID are listed in S4 Table.

The process encompassed the following steps:

1. an annotation-based draft network was generated using the PathoLogic program from the Pathway Tools suite, using the gbk file from the *C*. *crispus* genome annotation [25] and the metabolic reaction database MetaCyc20.5 [73].
2. an orthology-based network was generated using the protein sequences and metabolic network of *A*. *thaliana* (AraGEM, [57]), using the Pantograph software [74] to combine the output of ortholog searches with the Inparanoid and OrthoMCL softwares.
3. an orthology-based network was generated using the protein sequences from the well-annotated red microalga *Galdieria sulphuraria* [75] and its metabolic network reconstructed using Pathway Tools. This *G*. *sulphuraria* annotation-based network was then used as a template to generate a *C*. *crispus* network using Pantograph.
4. an orthology-based network was also generated using the protein sequences from the version 2 of the annotated genome of *E*. *siliculosus* [76], as well as version 2 of its metabolic network [55].
5. the four preliminary networks were merged together in the AuReMe environment, and an additional gap-filling step was performed using Meneco [77], constraining the network to produce the 84 metabolites from the literature that were indexed in the Metacyc database.

### Flux-balance analysis

A biomass reaction was established based on the previous *E*. *siliculosus* data [78]. One compound, L-alpha-alanine, gave negative fluxes, thus blocking biomass production. This was due to the absence of the alanine dehydrogenase reaction. The corresponding enzyme (CHC_T00008930001) was present in the *C*. *crispus* network but annotated as an NAD(P) transhydrogenase. We completed the annotation through the manual curation form to enable it to dehydrogenate alanine and to restore producibility of the biomass (http://gemaureme.irisa.fr/ccrgem/index.php/Manual-ala_dehy).

### Global metabolic networks comparisons

In order to compare the global features of the GSM from *C*. *crispus* with other ones, it is necessary to use the same reference database. This is the case for *E*. *siliculosus* and *E*. *subulatus* for which the reconstructions are based on MetaCyc [73] while *A*. *thaliana* and *C*. *reinhardtii* are respectively from KEGG [79] and BiGG [80]. To get access to MetaCyc pathway information for *A*. *thaliana* and *C*. *reinhardtii*, their networks were mapped using the sbml_mapping function implemented in the AuReMe workflow [55]. This function provides a dictionary of corresponding reactions from a database to an other one using the MetaNetX cross-reference database [81]. This dictionary was then used in AuReMe to create a new genome-scale metabolic network based on the new reference database for *A*. *thaliana* and *C*. *reinhardti*. Those new networks, who are comparable in size with the published ones (+/- 10 reactions and enzymes in our counts) enabled to estimate the number of pathways as defined in MetaCyc for both species.

### Ab-initio inference of new metabolic reactions

To enable the incorporation of the orphan metabolites that were not yet in MetaCyc into the network, we developed a new method called “Pathmodel” that can infer new reactions based on molecular similarity and dissimilarity. This knowledge-based approach is founded on two modes of reasoning (deductive and analogical) and was implemented using a logic programming approach known as Answer Set Programming (ASP) [82, 83]. It is a declarative approach oriented toward combinatorial (optimization) problem-solving and knowledge processing. ASP combines both a high-level modeling language with high performance solving engines so that the focus is on the problem specification rather than the algorithmic part. ASP expresses a problem as a set of logical rules (clauses). Problem solutions appear as particular logical models (so-called stable models or answer sets) of this set. An ASP program consists of rules *h:- b_1_,…, b_m_ not b_m+1_*,…, *not b_n_*, where each *b*_*i*_ and *h* are literals and *not* stands for default negation. In fact, each proposition is a predicate, encoded by a function whose arguments can be constant atoms or variables over a finite domain. The rule states that the head *h* is proven to be true (*h* is in an answer set) if the body of the rule is satisfied, i.e. b_1_,…, bm are true and it cannot be proved that b_m+1_,…, bn are true.

In short, the main predicates used in Pathmodel to represent molecules and reactions forming a knowledge base are *bond, atom* and *reaction* on which several logical rules are then applied to all possible reactions and potential reactants. Resulting products that do not belong to the knowledge base but that correspond to an observed m/z ratio are considered as new inferred metabolites and reactions. The finally encoded reactions result from iterative interactions between analogical model construction, automated inference, and manual validation of inferred reactions with respect to experimental results. The principles of the encoded analogical reasoning are explained in the results and discussion part.

The source code is available in the following Gitlab repository: https://gitlab.inria.fr/DYLISS/PathModel. The added reactions are listed on the following pages: http://gem-aureme.irisa.fr/ccrgem/index.php/Manual-pathmodel_inference http://gem-aureme.irisa.fr/ccrgem/index.php/Manual-pathmodel_inference_new_rxn

### De novo gene prediction and manual curation of gene sequence models

Missing genes from the sterol synthesis pathway (squalene monooxygenase and sterol C-4 methyl oxidase) were found by targeted tblastn using orthologs from other organisms as a query. The new gene predictions are provided in supplementary dataset 1 and will be included in the next version of *Chondrus crispus* genome browser (http://mmo.sb-roscoff.fr/jbrowse/?data=data%2Fpublic%2Fchondrus). The split protein sequence of sterol delta-7 reductase was also restored as a single protein prediction, merging the two adjacent partial predictions.

### Phylogenetic analyses

Collected sequences were aligned using Clustal Omega [84] and alignments were checked manually and edited with Seaview [85]. Phylogenetic trees were built using PHYML [86] using the LG model [87] with a gamma law. The reliability of nodes was assessed by likelihood-ratio test [88].

## Acknowledgments

We thank Cécile Hervé for help in collecting field samples from *C*. *crispus*, Gaëlle Correc for help in sample preparation, Karine Cahier for help during GC-MS analyses on the MetaboMer-Corsaire plateform, Jeanne Got and Marie Chevallier for help in using preliminary versions of AuReMe, and Clémence Frioux for help in analyzing FBA artefacts. GVM is also grateful to Ralf J. Sommer for giving him the possibility to start developping Pathmodel during a previous postdoctoral stay at the Max-Planck Institute for Developmental Biology in Tübingen.

## Author contributions

Conceptualization: GVM, LD, SMD, PS, EC, JN, CB, CL, AS, JC

Data curation: GVM, JG, MA, AB, CL, LD, SMD, PS, EC, CB, CL, AS, JC

Funding acquisition: AS, SMD, CB, CL

Investigation: GVM, JG, MA, AB, CT, CM

Project administration: GVM

Software & Methodology: AB, GVM, JN

Writing – original draft: AB, JG, MA, PS, JN, AS, GVM

Writing – review & editing: AB, JG, LD, SMD, PS, CT, EC, JN, CL, CB, JC, AS, GVM

