## Supplementary material for "Inferring biochemical reactions and metabolite structures to cope with metabolic pathway drift": S3-S4 Tables

### Table S3

| Usual name | Category | MetaCyC ID | References |
|---|---|---|---|
| dodecanoic acid | 12:0 fatty acid | DODECANOATE | Santos et al., 2015 ; Robertson et al., 2015 |
| Myristic acid | 14:0 fatty acid | CPD-7836 | Pettitt et al., 1989; Tasende, 2000; Van Ginneken et al., 2011; Robertson et al., 2015 ; Belghit et al., 2017 |
| Pentadecanoic acid | 15:0 fatty acid | CPD-8462 | Santos et al., 2015. Belghit et al., 2017 |
| Palmitic acid | 16:0 fatty acid | PALMITATE | Pettitt et al., 1989; Tasende, 2000; Van Ginneken et al., 2011; Robertson et al., 2015 ; Belghit et al., 2017 |
| Heptadecanoic acid | 17:0 fatty acid | CPD-7830 | Santos et al., 2015 |
| Stearic acid | 18:0 fatty acid | STEARIC_ACID | Tasende et al., 2000 ; Robertson et al., 2015 |
| Eicosanoic acid | 20:0 fatty acid | ARACHIDIC_ACID | Santos et al., 2015 |
| Docosanoic acid | 22:0 fatty acid | DOCOSANOATE | Santos et al., 2015 |
| Tricosanoic acid | 23:0 fatty acid | CPD-7834 | Santos et al., 2015 |
| Tetracosanoic acid | 24:0 fatty acid | TETRACOSANOATE | Santos et al., 2015 |
| Palmitoleic acid | 16:1(n-7) fatty acid | CPD-9245 | Pettitt et al., 1989; Tasende, 2000; Robertson et al., 2015 ; Belghit et al., 2017 |
| Oleic acid | 18:1(n-9) fatty acid | OLEATE-CPD | Tasende et al., 2000; Van Ginneken et al., 2011; Robertson et al., 2015 ; Belghit et al., 2017 |
| Linoleic acid | 18:2(n-6) fatty acid | LINOLEIC_ACID | Tasende et al., 2000 ; Robertson et al., 2015 ; Belghit et al., 2017 |
| Alpha Linolenic acid | 18:3(n-3) fatty acid | LINOLENIC_ACID | Tasende et al., 2000 |
| γ-linolenic acid | 18:3(n-6) fatty acid | CPD-8117 | Robertson et al., 2015 ; Belghit et al., 2017 |
| Octadecatetraenoic acid | 18:4(n-3) fatty acid | CPD-12653 | Tasende et al., 2000 ; Robertson et al., 2015 ; Belghit et al., 2017 |
| Arachidonic acid | 20:4(n-6) fatty acid | ARACHIDONIC_ACID | Tasende et al., 2000 ; Banskota et al., 2014 ; Robertson et al., 2015 ; Belghit et al., 2017 |
| Eicosapentaenoic acid | 20:5(n-3) fatty acid | 5Z8Z11Z14Z17Z-EICOSAPENTAENOATE | Tasende et al., 2000 ; Banskota et al., 2014 ; Robertson et al., 2015 ; Belghit et al., 2017 |
| Octanedioic acid | fatty acid | CPD0-1264 | Santos et al., 2015 |
| Nonanedioic acid | fatty acid | CPD0-1265 | Santos et al., 2015 |
| Cycloartenol | sterol | CYCLOARTENOL | Saito and Idler, 1966; Alcaide et al., 1968 |
| Cholesterol | sterol | CHOLESTEROL | Saito and Idler, 1966; Tasende et al., 2000 ; Santos et al., 2015 |
| 7-Dehydrocholesterol | sterol | 7-DEHYDROCHOLESTEROL | Tasende et al., 2000 |
| Brassicasterol | sterol | BRASSICASTEROL | Saito and Idler, 1966 ; Tasende et al., 2000 |
| Campesterol | sterol | CAMPESTEROL | Tasende et al., 2000 ; Santos et al., 2015 |
| 24-Methylenecholesterol | sterol | 24-METHYLENECHOLESTEROL | Tasende et al., 2000 |
| Sitosterol | sterol | SITOSTEROL | Saito and Idler, 1966; Tasende et al., 2000 ; Santos et al., 2015 |
| Stigmasterol | sterol | STIGMASTEROL | Tasende et al., 2000 |
| 15-keto-prostaglandin E2 | oxylipin | HYDROXY-915-DIOXOPROSTA-13-ENOATE | Gaquerel et al., 2007 |
| lutein | carotenoid | LUTEIN | Banskota et al., 2014 |
| Chlorophyll a | tetrapyrrole | CHLOROPHYLL-A | Melo et al., 2015 ; Robertson et al., 2015 |
| all-trans-beta-carotene | carotenoid | CPD1F-129 | Robertson et al., 2015 |
| 9-cis-betacarotene | carotenoid | CPD-14646 | Robertson et al., 2015 ; Belghit et al., 2017 |
| zeaxanthin | carotenoid | CPD1F-130 | Robertson et al., 2015 |
| 2,6,6-trimethyl-1,3-cyclohexadiene-1-carboxaldehyde (safranal) | carotenoid | CPD-8669 | Pina et al., 2014 |
| Alanine | aminoacid | L-ALPHA-ALANINE | Young et al., 1958, Belghit et al., 2017 |
| Arginine | aminoacid | ARG | Young et al., 1958, Belghit et al., 2017 |
| Aspartic acid | aminoacid | L-ASPARTATE | Young et al., 1958, Belghit et al., 2017 |
| Citrulline | aminoacid | L-CITRULLINE | Young et al., 1958 ; Belghit et al., 2017 |
| Cystine | aminoacid | CYSTINE | Young et al., 1958 |
| Glutamic acid | aminoacid | GLT | Young et al., 1958 ; Belghit et al., 2017 |
| Glycine | aminoacid | GLY | Young et al., 1958 ; Belghit et al., 2017 |
| Histidine | aminoacid | HIS | Young et al., 1958 ; Belghit et al., 2017 |
| Isoleucine | aminoacid | ILE | Young et al., 1958 ; Belghit et al., 2017 |
| Leucine | aminoacid | LEU | Young et al., 1958 ; Belghit et al., 2017 |
| Lysine | aminoacid | LYS | Young et al., 1958 ; Belghit et al., 2017 |
| Methionine | aminoacid | MET | Young et al., 1958 ; Belghit et al., 2017 |
| Ornithine | aminoacid | L-ORNITHINE | Young et al., 1958 ; Belghit et al., 2017 |
| Phenylalanine | aminoacid | PHE | Young et al., 1958 |
| Proline | aminoacid | PRO | Young et al., 1958 ; Belghit et al., 2017 |
| Serine | aminoacid | SER | Young et al., 1958 ; Belghit et al., 2017 |
| Threonine | aminoacid | THR | Young et al., 1958 ; Belghit et al., 2017 |
| Tyrosine | aminoacid | TYR | Young et al., 1958 ; Belghit et al., 2017 |

### Table S3

| | | | |
|---|---|---|---|
| Valine | aminoacid | VAL | Young et al., 1958 ; Belghit et al., 2017 |
| Shinorine | Mycosporine-like aminoacid | CPD-18778 | Kräbs et al., 2004 |
| UDP-α-D-galactose | nucleotide sugar | CPD-14553 | Collén et al., 2014 |
| ι-carrageenose | carrageenan | Iota-Carrageenan | Matsuhiro et al., 1992 |
| ν-carrageenan | carrageenan | Nu-Carrageenan | Matsuhiro et al., 1992 |
| Glycerol | polyol | GLYCEROL | Santos et al., 2015 |
| Heptadecane | alcane | HEPTADECANE-CPD | Santos et al., 2015 |
| 6,10,14-Trimethyl-2-pentadecanone | methylketone | CPD-7875 | Santos et al., 2015 |
| Hexadecan-1-ol | Long chain aliphatic alcohol | CPD-348 | Santos et al., 2015 |
| 9-Octadecen-1-ol | Long chain aliphatic alcohol | CPD-7873 | Santos et al., 2015 |
| Docosan-1-ol | Long chain aliphatic alcohol | CPD-7845 | Santos et al., 2015 |
| Octacosan-1-ol | Long chain aliphatic alcohol | CPD-7872 | Santos et al., 2015 |
| acetaldehyde | aldehyde | ACETALD | Pina et al., 2014 |
| 2-methypropanal | aldehyde | BUTANAL | Pina et al., 2014 |
| Butanal | aldehyde | CPD-7031 | Pina et al., 2014 |
| 3-methybutanal | aldehyde | METHYLBUT-CPD | Pina et al., 2014 |
| Pentanal | aldehyde | CPD-9053 | Pina et al., 2014 |
| Hexanal | aldehyde | HEXANAL | Pina et al., 2014 |
| Benzaldehyde | aldehyde | BENZALDEHYDE | Pina et al., 2014 |
| Ethanol | short chain aliphatic alcohol | ETOH | Pina et al., 2014 |
| 1-butanol | short chain aliphatic alcohol | BUTANOL | Pina et al., 2014 |
| 1-pentanol | short chain aliphatic alcohol | PENTANOL | Pina et al., 2014 |
| 2-butanone | short chain ketone | ACETONE | Pina et al., 2014 |
| 3,5-octadien-2-one | short chain ketone | MEK | Pina et al., 2014 |
| dichloromethane | halocarbon | CPD-681 | Pina et al., 2014 |
| chloroform | halocarbon | CPD-843 | Pina et al., 2014 |
| 2-methylpropanoic acid | carboxylic acid | ACET | Pina et al., 2014 |
| 2-methylbutanoic acid | carboxylic acid | ISOBUTYRATE | Pina et al., 2014 |
| hexane | alcane | CPD-9288 | Pina et al., 2014 |
| 2,2,4-trimethylpentane | alcane | CPD-19039 | Pina et al., 2014 |

#### Table S4

| Usual name | Category | References |
|---|---|---|
| Heneicosanoic acid | 21:0 fatty acid | Santos et al., 2015 |
| N/A | 15:1 fatty acid | Robertson et al., 2015 |
| N/A | 18:1(n-7) fatty acid | Robertson et al., 2015 |
| 10-nonadecenoate | 19:1(n-9) fatty acid | Belghit et al., 2017 |
| Eicosadienoic acid | 20:2(n-6) fatty acid | Robertson et al., 2015 |
| Eicosatrienoic acid | 20:3(n-6) fatty acid | Robertson et al., 2015 |
| Docosadienoate | 22:2(n-6) fatty acid | Belghit et al., 2017 |
| Octadeca-9-enoic acid | fatty acid | Santos et al., 2015 |
| 22-Dehydrocholesterol | sterol | Tasende et al., 2000 |
| 11-hydroxy-octadecadienoic acid (11-HODE) | oxylipin | Gaquerel et al., 2007 |
| 13-hydroxy-9Z,11E-octadecadienoic acid (13-HODE) | oxylipin | Gaquerel et al., 2007; Belghit et al., 2017 |
| 13S-hydroxy-9Z,11E,15Z-octadecatrienoic acid (13-HOTrE) | oxylipin | Belghit et al., 2017 |
| 13-oxo-9Z,11E-octadecadienoic acid (13-oxo-ODE) | oxylipin | Gaquerel et al., 2007 |
| 13-hydroxyeicosatrienoic acid (13-HETrE) | oxylipin | Gaquerel et al., 2007 |
| 13-hydroxyeicosatetraenoic acid (13-HETE) | oxylipin | Gaquerel et al., 2007 |
| 13-hydroxyeicosapentaenoic acid (13-HEPE) | oxylipin | Gaquerel et al., 2007 |
| 15-hydroxydocosahexaenoic acid (15-HDHE) | oxylipin | Gaquerel et al., 2007 |
| 11-hydroxyoctadecadienoic acid (11-HETE) | oxylipin | Gaquerel et al., 2007 |
| Hydroxypheophytin a | tetrapyrrole | Melo et al., 2015 |
| Pheophytin d | tetrapyrrole | Melo et al., 2015 |
| Hydroxypheophytin d | tetrapyrrole | Melo et al., 2015 |
| Monogalactosyldiacylglycerol 2 (MGDG2) | galactolipid | Pettitt et al., 1989 |
| Digalactosyldiacylglycerols (DGDG) | galactolipid | Pettitt et al., 1989 |
| Sulfoquinovosyldiacylglycerol 1 (SQDG1) | galactolipid | Pettitt et al., 1989 |
| Sulfoquinovosyldiacylglycerol 2 (QDG2) | galactolipid | Pettitt et al., 1989 |
| (2S)-1,2-bis-O-eicosapentaenoyl-3-O-β-D-galactopyranosylglycerol | galactolipid | Banskota et al., 2014 |
| (2S )-1-O-eicosapentaenoyl-2-O-arachidonoyl-3-O-β-D-galactopyranosylglycerol | galactolipid | Banskota et al., 2014 |
| (2S )-1-O -(6Z,9Z,12Z,15Z- octadecatetranoyl)-2-O-palmitoyl-3-O-β-D – galactopyranosylglycerol | galactolipid | Banskota et al., 2014 |
| (2S )-1-O -eicosapentaenoyl-2- O -palmitoyl-3-O -β -D -galactopyranosylglycerol | galactolipid | Banskota et al., 2014 |
| (2S )-1, 2-bis -O -arachidonoyl-3-O -β -D -galactopyranosylglycerol | galactolipid | Banskota et al., 2014 |
| (2S )-1-O -arachidonoyl-2-O -palmitoyl-3-O -β -D – galactopyranosylglycerol | galactolipid | Banskota et al., 2014 |
| (2S )-1-O-eicosapentaenoyl-2-O-palmitoyl-3-O-(β-D-galactopyranosyl-6-1α-D–galactopyranosyl)-glycerol | galactolipid | Banskota et al., 2014 |
| (2S)-1-O-arachidonoyl-2-O-palmitoyl-3-O-(β-D-galactopyranosyl-6-1α-D-galactopyranosyl)-glycerol | galactolipid | Banskota et al., 2014 |
| diphosphatidylglycerol, phosphatidic acid | DPG/PA | Pettitt et al., 1989 |
| l-citrullinyl-l-arginine | aminoacid | Laycock et al., 1977 |
| Gigartinine | aminoacid | Laycock et al., 1977 |
| Amide N | aminoacid | Young et al., 1958 |
| Asterina-330 | Mycosporine-like aminoacid | Athukorala et al., 2016; Guihéneuf et al., 2018 |
| MAA1 | Mycosporine-like aminoacid | This study |
| MAA2 | Mycosporine-like aminoacid | This study |
| Palythine | Mycosporine-like aminoacid | Karsten et al., 1998; Athukorala et al., 2016; Guihéneuf et al., 2018 |
| Palythene | Mycosporine-like aminoacid | Karsten et al., 1998 |
| Palythinol | Mycosporine-like aminoacid | Karsten et al., 1998; Athukorala et al., 2016 |
| Isofloridoside | heteroside | Kremer et al., 1982 |
| 1-Monohexadecanoin | Long chain aliphatic alcohol | Santos et al., 2015 |
| Tetradecan-1-ol | Long chain aliphatic alcohol | Santos et al., 2015 |
| Octadecan-1-ol | Long chain aliphatic alcohol | Santos et al., 2015 |
| 1-penten-3-ol | short chain aliphatic alcohol | Pina et al., 2014 |
| 2(Z)-penten-1ol | short chain aliphatic alcohol | Pina et al., 2014 |
| 3-methylbutanoic acid | carboxylic acid | Pina et al., 2014 |
| 2-methylbutanal | aldehyde | Pina et al., 2014 |
| 1-octen-3-ol | short chain aliphatic alcohol | Pina et al., 2014 |

| | | |
|---|---|---|
| 2,6-dimethylpyrazine | short chain ketone | Pina et al., 2014 |
| 2-propanone | short chain ketone | Pina et al., 2014 |
| acetic acid, anhydride | carboxylic acid | Pina et al., 2014 |
| 2,2,3-trimethylpentane | alcane | Pina et al., 2014 |
| tetradecane | alcane | Pina et al., 2014 |
