## Supplementary material for "Inferring biochemical reactions and metabolite structures to cope with metabolic pathway drift": S1-S5 Figs, S1-S2 Tables

**This file includes:**

Figures S1 to S6
Table S1 to S2
Supplementary dataset

**Other supplementary material for this manuscript includes the following:**

Tables S3 to S4 (separate pdf, content will be submitted to the Metabolights database)

| **Analysed compounds** | **Molecular weight** (g.mol-1) | **RT** (min) | **m/z $[M+H]^+$** (TMS) |
|---|---|---|---|
| brassicasterol | 398.66 | 25.5 | 470 |
| campesterol | 400.68 | 26.6 | 472 |
| 5α-cholestane | 372.67 | 20.3 | 372 |
| cholesterol | 386.65 | 24.7 | 458 |
| cycloartanol | 428.75 | 30.5 | 500 |
| cycloartenol | 426.72 | 29.0 | 498 |
| cycloeucalenol | 426.73 | 30.4 | 498 |
| 7-dehydrocholesterol | 384.63 | 25.6 | 456 |
| desmosterol | 384.64 | 25.5 | 456 |
| ergosterol | 396.65 | 26.2 | 468 |
| fucosterol | 412.69 | 28.2 | 484 |
| lanosterol | 426.39 | 27.8 | 498 |
| lathosterol | 386.65 | 25.8 | 458 |
| β-sitosterol | 414.39 | 28.0 | 486 |
| squalene | 410.72 | 19.8 | 482 |
| stigmasterol | 412.69 | 27.0 | 484 |
| Zymosterol | 384.64 | 25.9 | 456 |

**Supp. Table 3. Retention times and m/z ratio for analytical standards of sterols on a 7890-5975C Agilent GC-MS.**

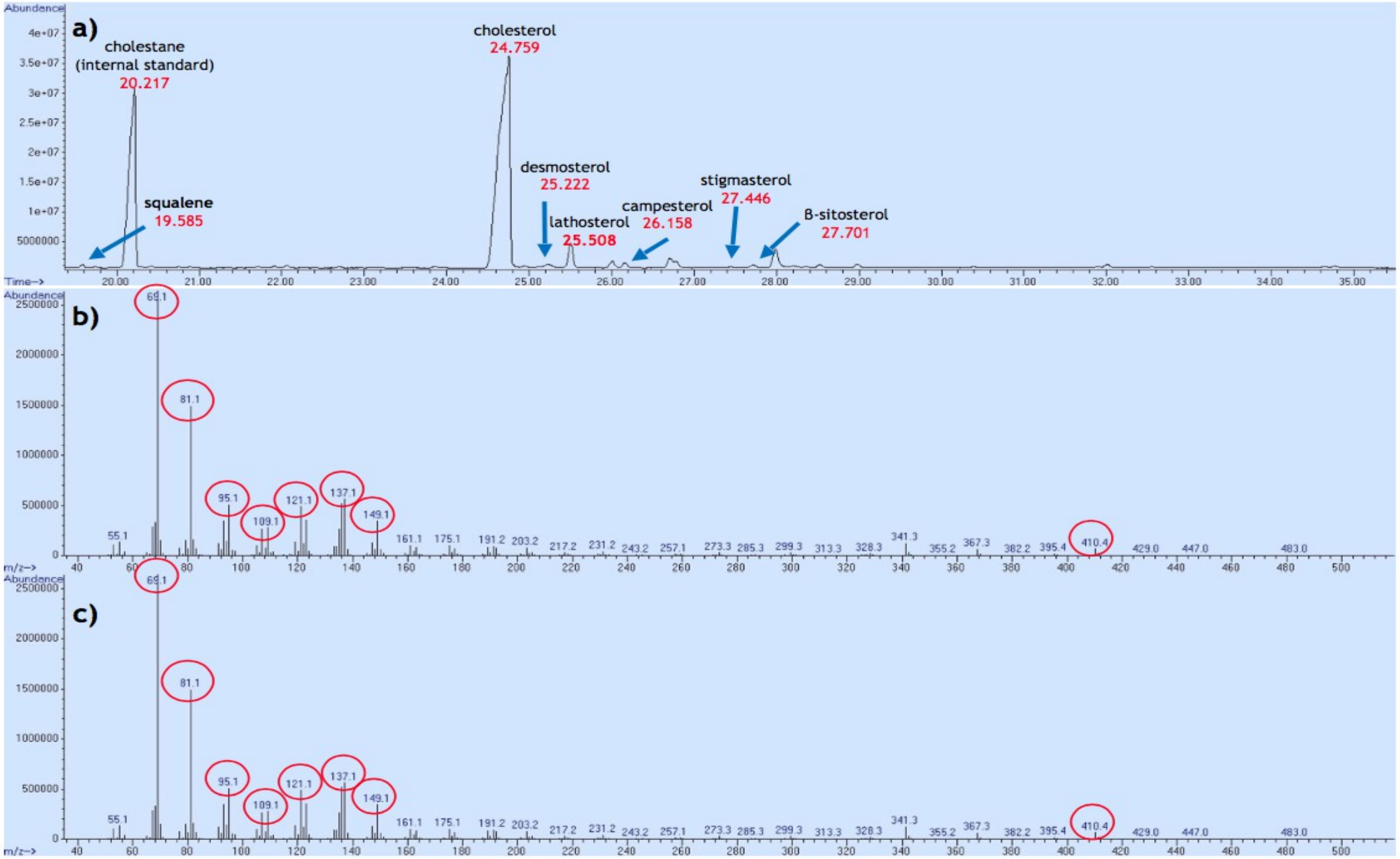


**Supp. Figure 1. Identification of squalene in *C. crispus*.** a) Total Ion Chromatogramm (TIC) from *C. crispus* extract. b) MS spectrum of squalene in *C. crispus* extract. c) MS spectrum of the squalene analytical standard. Main fragmentation peaks identical in both spectra are highlighted in red circles.

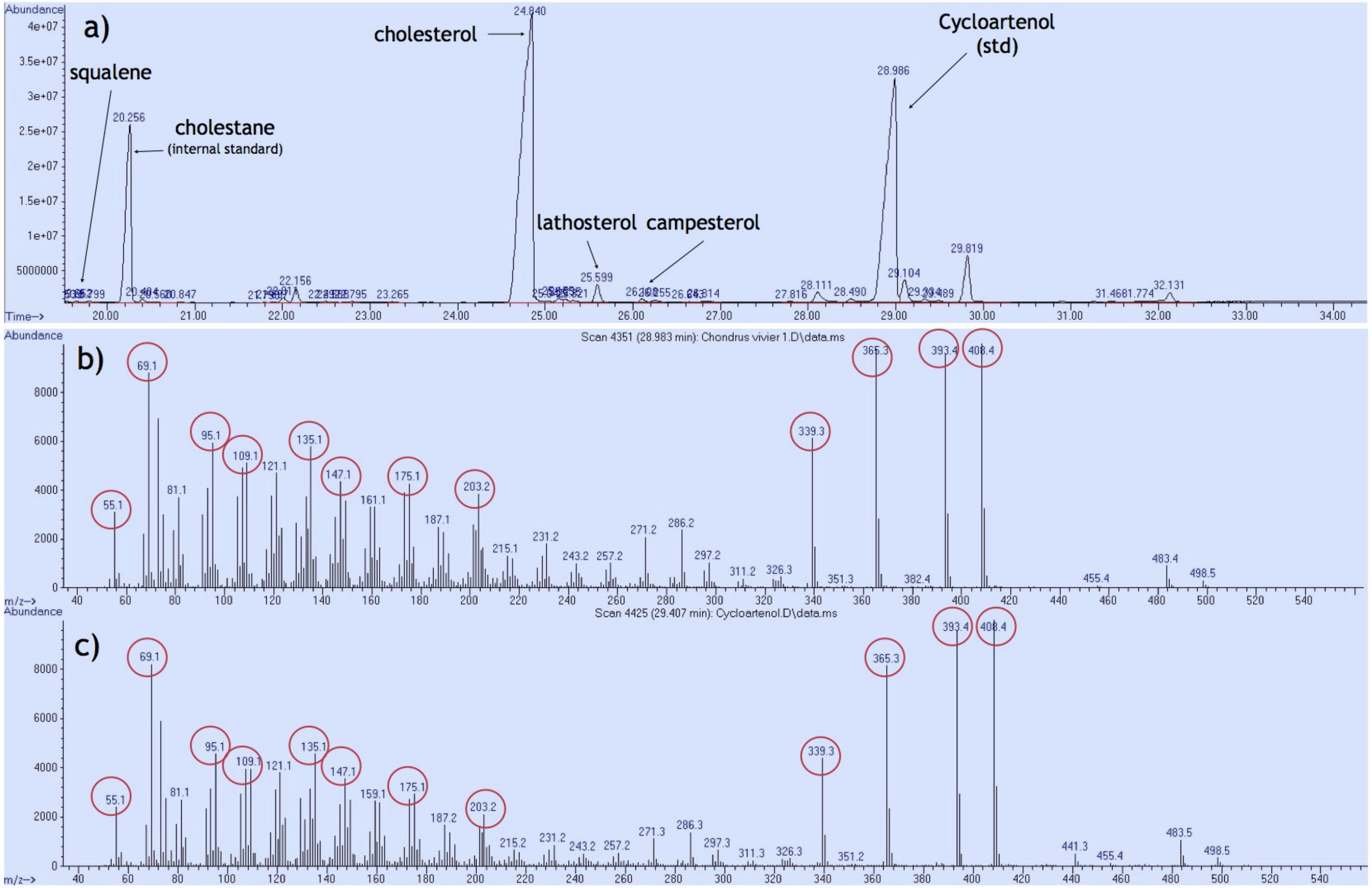


**Supp Figure 2. Control for technical detectability of cycloartenol in spiked *Chondrus crispus* extract.**
a) TIC from *Chondrus crispus* extract incubated with cycloartenol. b) MS spectrum of cycloartenol standard incorporated in *C. crispus* extract. c) MS spectrum of cycloartenol standard alone. Main fragmentation peaks identical in both spectra are highlighted in red circles.

| MAAs | Palythine | Mycosporine-glycine | MAA1 | Isujirene/Palythene | Asterina-330 | Palythinol or MAA2 | Shinorine | Porphyra-334 |
|---|---|---|---|---|---|---|---|---|
| Rt (min.) | 8.3 | 20.0 | 10.8 | 19.3 | 8.7 | 10.1 | 18.5 | 19.5 |
| m/z [M+H]+ observed | 245.1090 | 246.0932 | 271.1241 | 285.1401 | 289.1349 | 303.1497 | 333.1245 | 347.1399 |
| m/z calculated | 245.1132 | 246.0972 | 271.1288 | 285.1445 | 289.1394 | 303.1551 | 333.1292 | 347.1449 |
| EIC (Intens. x108) | | | | | | | | |
| *C. crispus* (April) | 16118542 | 707375 | 3254803 | 209911 | 5116637 | 26945 | 3129533 | 353130 |
| *C. crispus* (July) | 12600749 | 85700 | 928894 | 36714 | 3788544 | 18560 | 394887 | 11021 |
| *C. crispus* (August) | 16469850 | 219296 | 857212 | 238033 | 5653618 | 32998 | 1063642 | 83569 |
| *C. crispus* (Sept.) | 11230824 | 56477 | 2546286 | 77636 | 2917730 | < LOD | 5199737 | 33580 |
| UV (mAU) | | | | | | | | |
| *C. crispus* (April) | 20420 | 31,525 | 1541 | < LOD | 6996 | < LOD | 2299 | 117 |
| *C. crispus* (July) | 14578 | 171,83 | 1487 | < LOD | 7106 | 327,43 | 335 | < LOD |
| *C. crispus* (August) | 19005 | 242,7 | 2143 | < LOD | 9927 | 707,57 | 1245 | 248 |
| *C. crispus* (Sept.) | 12768 | < LOD | 989 | < LOD | 6367 | < LOD | 5128 | 136 |

**Supp. Table 4. MAAs composition in *Chondrus crispus* by LC-UV-HRMS. Extracted Ion Chromatogramm (EIC) of selected MMAs were obtained in positive mode; UV Absorbance was recorded at 330 nm (LOD = Limit Of Detection).**

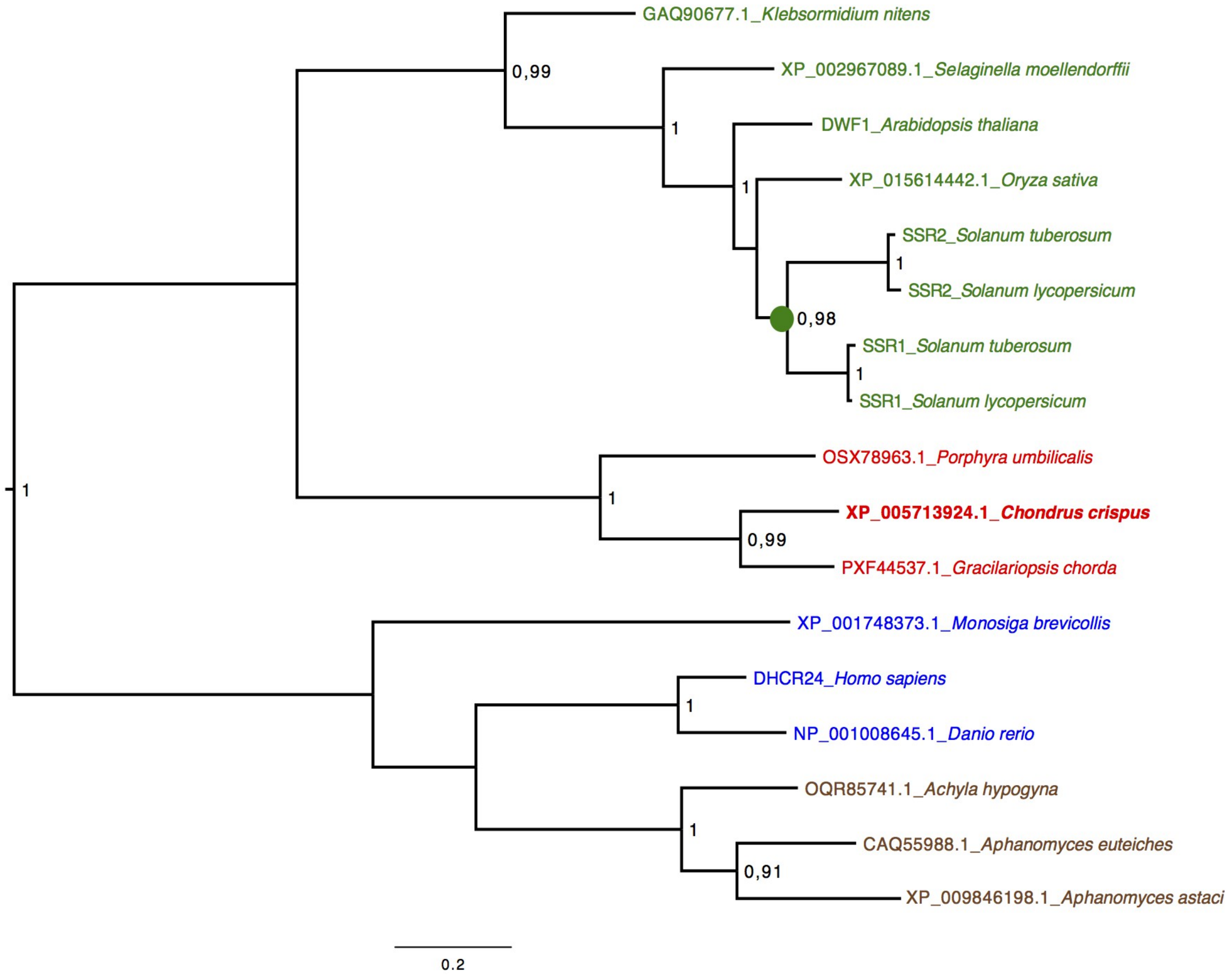


**Supp. Figure 3. Maximum-likelihood tree of eukaryotic side-chain reductases.** In green: sequences from green plants (streptophytes). The green dot indicates lineage-specific duplication in solanaceans. In red: sequences from red algae. In blue: sequences from opistokonts (vertebrates + choanoflagellates). In brown: sequences from oomycete stramenopiles. Likelihood-ratio test values above 0.90 are indicated. Those above 0.97 are considered significant.

**Supplementary dataset S1.** New or edited protein sequences for the sterol synthesis pathway in *Chondrus crispus*.

>scaffold90:7511-6165(-) candidate squalene epoxidase
RDGRRVLCVERQLYAPSGALCAPPRIVGELLQPGGYDALCRLGLADALLDIDAQVIRGYA
LFLGPRAERLPYHQPGGPDPDPDPAARPQPEGRAFHNGRFLKRLREIARAHPNV
TLVEGNVLALLERDGAVVGVRYATRGNKAATAHAGLTIAC
DGCGSALRKRAAAHHHVTVYSNFHGLVLHVPALPFPNHGHVVLADPCPVLFYPISATEVR
CLVDIPSTYAGDAAEYILHTVVPQVPPPLRAPLATAVRERRSKMMPNRVMPAPA
HVVPGAVLLGDAFNMRHPLTGGGMTVALTDVELLRELLAPVPDLSDAPAVAAKLQLFYER
RKPMSTTINILANALYTLFCATDDPALRDMRAACLDYLAKGGRMTHDPIAMLGGLKPQRH
LLLAHFFAVALYGCGKALMPFPTPARLVRAWSIFRASFNIIKPLANAEGFWPLSWLPLNSL

>scaffold20:461442-460650(-) candidate squalene epoxidase
LCRLGLADALLHIDAQVIRGYALFLGPRAERLPYHQPGEPDPDPAARPQPEG
RAFHNGRFLKRLREIARAHPNVTLIEGNVLALLERDGAVV
GVRYATRGNKAATAHAGLTIACDGCGSALRKRAAAHHHVTVYSNFHGLVLHVPALPFPNH
GHVVLAHPCPVLFYPISATEVRCLVDL
YILHTVVPQVPPSLRAPLATTVRERRSKMMPNRVMPAPAHVVPGAVLLGDAFNMRHPLTG
GGMTVALTDVELLRGLLAP

>scaffold57:152407-364140(+) candidate squalene epoxidase
RFAGPEHPSCGLKPQRHLLLAHFFAVALYGCGKALMPFPTPARPVRAWSIFRASFNFIK

PLANAEGFWPLSWLPLN

LCRLGLADALLDIDAQVIRGYALFLGPRAERLPY

LCRLGLADALLHIDAQVIRGYALFLGPRAERLPYHQPGGPDPDPAARPQPEG

RAFHNCRFLKRLREIARAHPNVTLIEGNVLALLERDGAVV

GVRYATRGNKAATAHAGLTIACDGCGSALRKRAAAHHHVTVYSNFHGLVLHVPALPFPNH

GHVVLAHPCPVLFYPISATEVRCLVDL

WSTYAGDAAEYILHTVVPQVPPSLRAPLATAVRERRSKMMPNRVMPAPAHVVPGAVLLGD

AFNMRHPLTGGGMTVALTDVELLRGLLAP

>scaffold212:177405-176674(-) candidate C-4 sterol methyl oxidase

WDLLCRHTRAYPMFVVGCFASQLAGYFLGCAPFVLLDALRARSTPFRKIQPGKYAPRRAV

FAAAAAMLRSFATVVLPLLAAGGLFIERVGISRDAPFPSPRVVLLQVAYFFLVEDFLNYW

VHRALHLPWLYTRVHSVHHEYDAPFAVVAAYAHPVEVVLQALPTFAGPLMLGPHLYTLCV

WQLFRNWEAIDIHSGYDHAWGLASVLPWYAGPEHHDFHHFLHSGNFASVFTWCDWAYGTD

LAYE

>CHC_T00006492-3001 fusion of adjacent protein predictions CHC_T00006492001 and CHC_T00006493001; candidate sterol delta-7 reductase

MLGIAAWKGFIRYGLLYDHFGEVLAFLGKFALVVTVLLYFRGIYFPTNSDSGTTSFGIVWDMWHGTELHP

EIFGVSLKQLVNCRFALMGWSVAIVAFACKQREQYGYVSNSMLVSVVLQLVYIFKFFVWEAGYFNSVSLD

HSHVCLFWIYLRPLY

MVGVGAICCNYWTDKQREVFRATNGQVTIWGQKPVSIEAQYVTGDGKKRRSLLLASGWWGVSRHVNYVFE

IALTFCWSVPAGGTGVIPYVYVMFLTILLTDRAYRDEVRCSEKYGKYYEEYCRLVPYKMIPGVY

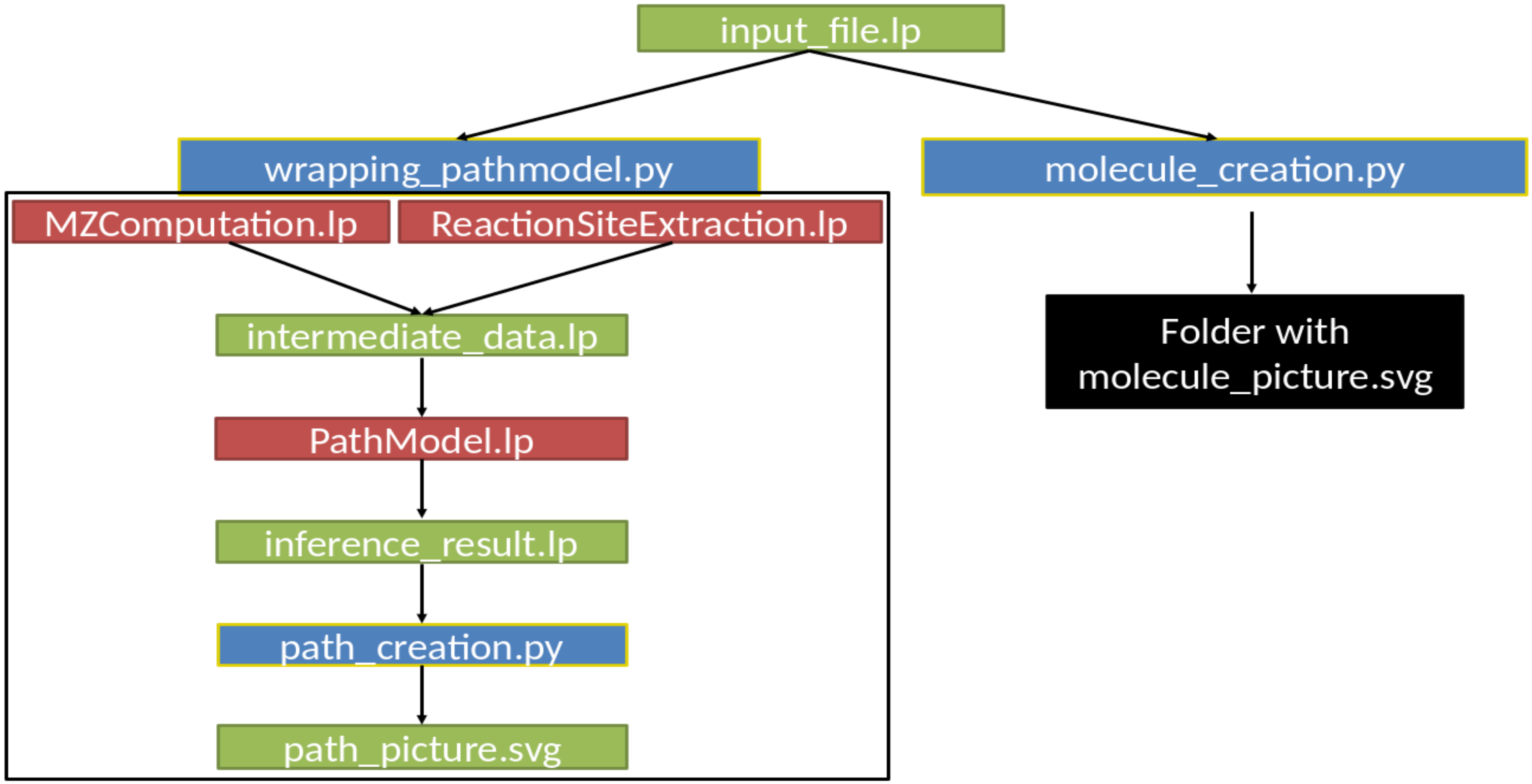


**Supp. Figure 4. Architecture of Pathmodel scripts.** In green: Data files, either input or result files. In red: ASP scripts. In blue: Python scripts. In black: folder containing results from molecule_creation.py (molecule pictures). The black line shows the wrapping of all the scripts inside by wrapping_pathmodel.py.

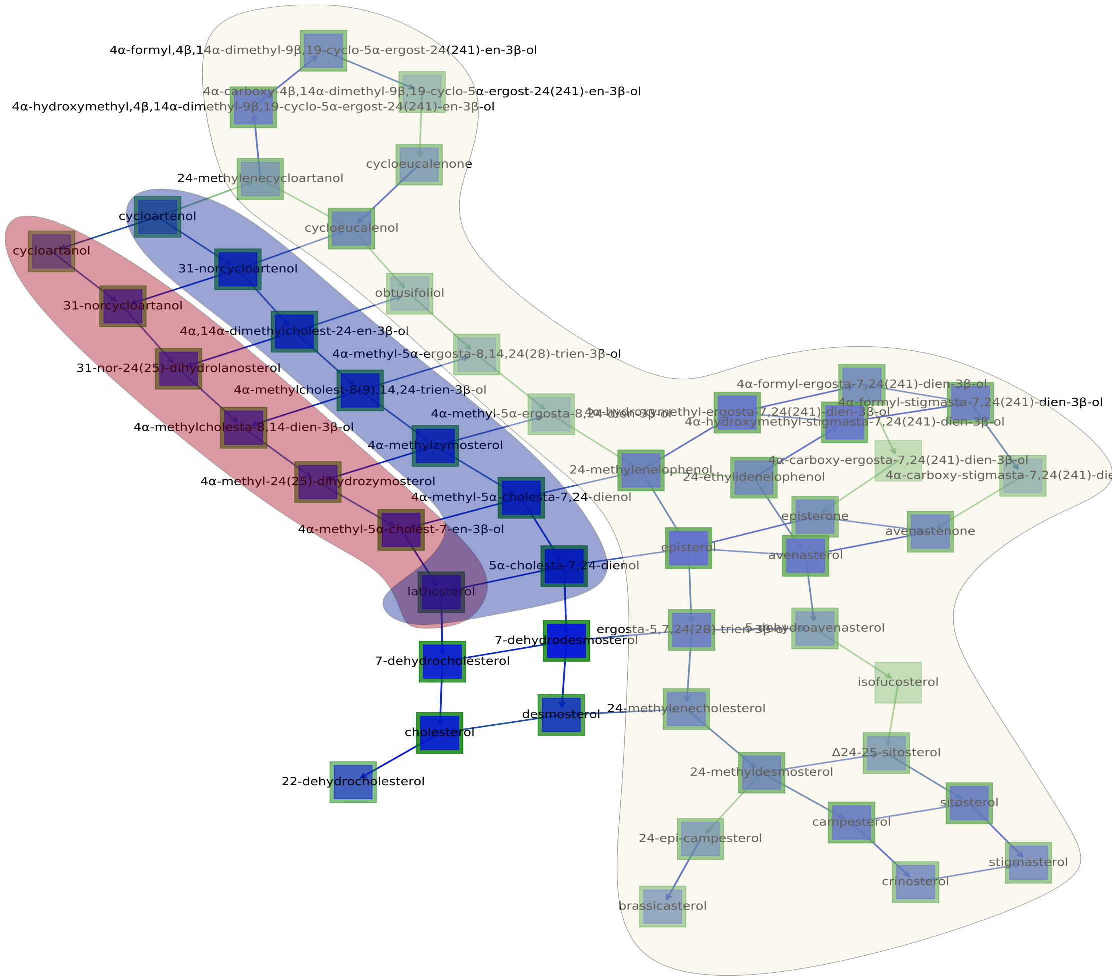


**Supp. Figure 5. Result of Pathmodel for sterol by path_creation.py.** In white: reactions and metabolites from PWY-2541 (Metacyc). In red: Early SSR pathways. In blue: Late SSR pathways. Arrows in green are from PWY-2541 and arrows in blue are inferred by Pathmodel.

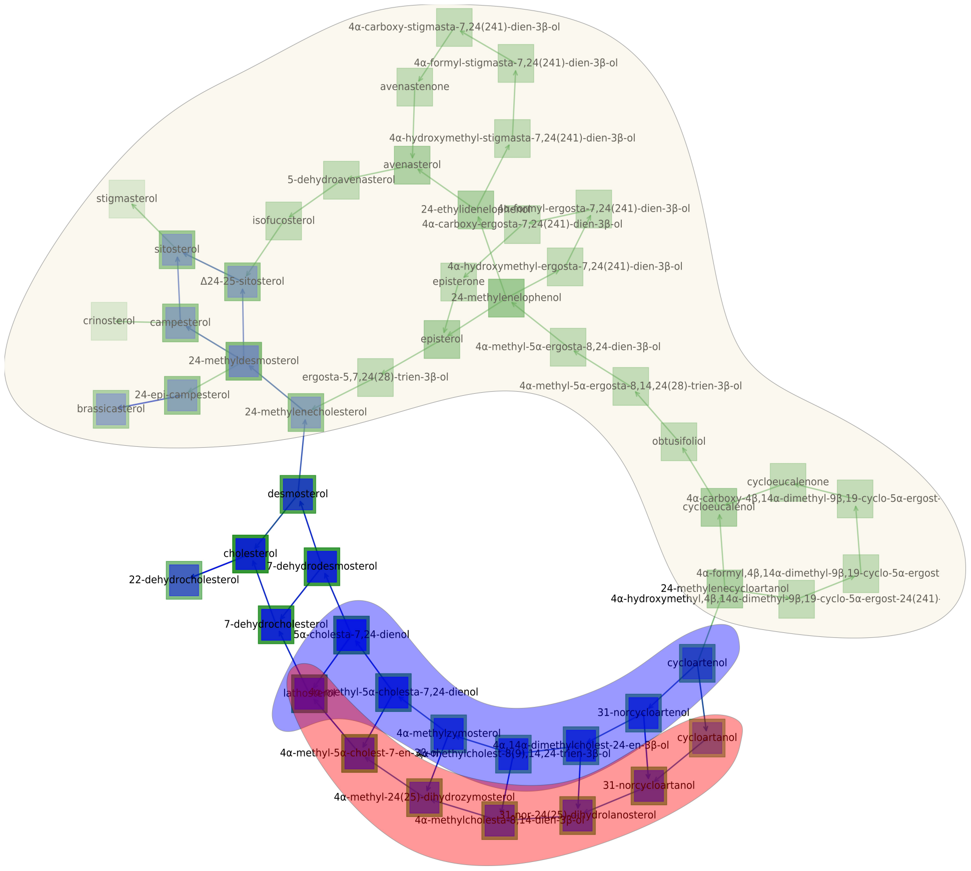


**Supp. Figure 6. Result of Pathmodel for sterol with path_creation.py and absent molecules.** In white: reactions and metabolites from PWY-2541 (Metacyc). In red: Early SSR pathways. In blue: Late SSR pathways. Arrows in green are from PWY-2541 and arrows in blue are inferred by Pathmodel. In this example, we put all the metabolites of the PWY-2541 as absent molecules, so no inference was made on them.
